# Consistent consensus-based annotation of spatial adaptive immune receptor repertoires from long-read sequencing using *LongAIRR*

**DOI:** 10.64898/2026.06.22.733709

**Authors:** Jonas Schuck, Samira Ortega Iannazzo, Zeina Mahmoud, Charles Gwellem Anchang, Lucie-Marie Hasse, Katharina Weber, Katharina Imkeller

**Affiliations:** Goethe University, University Hospital, Institute of Neurology (Edinger Institute), Frankfurt, Germany; Frankfurt Cancer Institute (FCI), Frankfurt, Germany; University Cancer Center (UCT), Frankfurt, Germany

## Abstract

The combination of spatial transcriptomics with long-read sequencing enables spatial characterization of full-length transcripts within solid tissue sections. However, standardized computational analysis frameworks are lacking, and it remains unclear whether available long-read sequencing platforms from Oxford Nanopore Technologies and Pacific Biosciences yield comparable results.

Here, we present a computational strategy for spatial full-length transcript analysis, focusing on the spatial profiling of adaptive immune receptor repertoires (AIRR). Our approach introduces an adaptive filtering strategy that dynamically refines read selection and significantly improves consensus accuracy, enabling high-confidence sequence reconstruction independent of platform-specific sequencing error profiles. We further derive evidence-based guidelines tailored to the consistent and robust analysis of spatial AIRR data.

The resulting software *LongAIRR* is modular and interoperable with existing spatial transcriptomics and AIRR analysis frameworks. This work establishes a methodological foundation for spatial immunology, enabling precise mapping of immune repertoires within their native tissue microenvironments.

## INTRODUCTION

Spatial transcriptomics technologies enable sequencing and quantification of transcripts in their spatial context and provide a detailed characterization of cellular heterogeneity in solid tissues (Ståhl2016). Platforms such as 10x Genomics Visium V1, and Visium HD 3′ directly capture and spatially barcode mRNA molecules, enabling the assessment of spatially resolved transcript heterogeneity. When combined with long-read sequencing technologies such as Oxford Nanopore Technologies (ONT) or Pacific Biosciences (PacBio), these approaches allow precise spatial characterization of full-length transcript characteristics, including isoform usage, splice variants and other structural features (Amarasinghe2020, Logsdon2020).

A key application of full-length spatial transcript sequencing is the mapping of adaptive immune receptor repertoires (AIRR) in solid tissues (Benotmane2023, Engblom2023, Guo2025, Hudson2022, Liu2022, Ly2025, Meylan2022, Sudmeier2022). B cell receptors (BCRs) and T cell receptors (TCRs) are surface-expressed molecules that enable adaptive immune cells to recognize their cognate antigens. During lymphocyte development, the transcripts encoding BCR chains (Ig γ1-γ4, Ig α1-α2, Ig μ, Ig ε, Ig δ, Ig κ, Ig λ) and TCR chains (TCR α, TCR β, TCR γ, TCR δ) undergo V(D)J recombination, a process in which variable (V), diversity (D), and joining (J) gene segments are rearranged to generate a unique antigen receptor sequence in each developing B or T cell (Schatz2011). Determining antigen receptor sequences at spatial resolution enables tracking of adaptive immune responses within solid tissue. For example, it allows detection of clonal lymphocyte expansion and, in B cells, mapping of somatic hypermutation and isotype class switching during affinity maturation in adaptive immune responses. Spatial adaptive immune receptor repertoire (spAIRR) data therefore provides insights into antigen specificity and clonal relationships of B and T cells in the tissue and has become an increasingly powerful tool for studying immune responses in infection, autoimmune disease, or cancer (Engblom2023, Liu2022, Meylan2022, Sudmeier2022).

Several experimental approaches have recently been developed to sequence spatial B and T cell repertoires (Benotmane2023, Engblom2023, Hudson2022, Jiang2026, Ly2025, Plumbom2025). These current state-of-the-art protocols employ probe-based enrichment or specific amplification of molecules from 10x Genomics Visium cDNA libraries, which contain spatially barcoded full-length cDNA molecules. In these workflows, a fresh-frozen tissue section is placed on a glass slide coated with spatially barcoded poly-A capture primers. The Visium V1 protocol captures transcripts at a spot-based resolution of 55 µm using primers containing a 16-nucleotide (nt) spatial barcode (SPBC) followed by a 12 nt unique molecular identifier (UMI). The more recent Visium HD 3′ protocol increases spatial resolution to a 2 µm grid, with capture primers containing an extended UMI-SPBC segment totaling 43 nucleotides (**Fig. 1a**). In both protocols, the SPBC and UMI are appended to the 3’ end of the poly(A)-tail-captured cDNA molecule and represent a unique molecular and spatial identifier.

**Figure 1:**
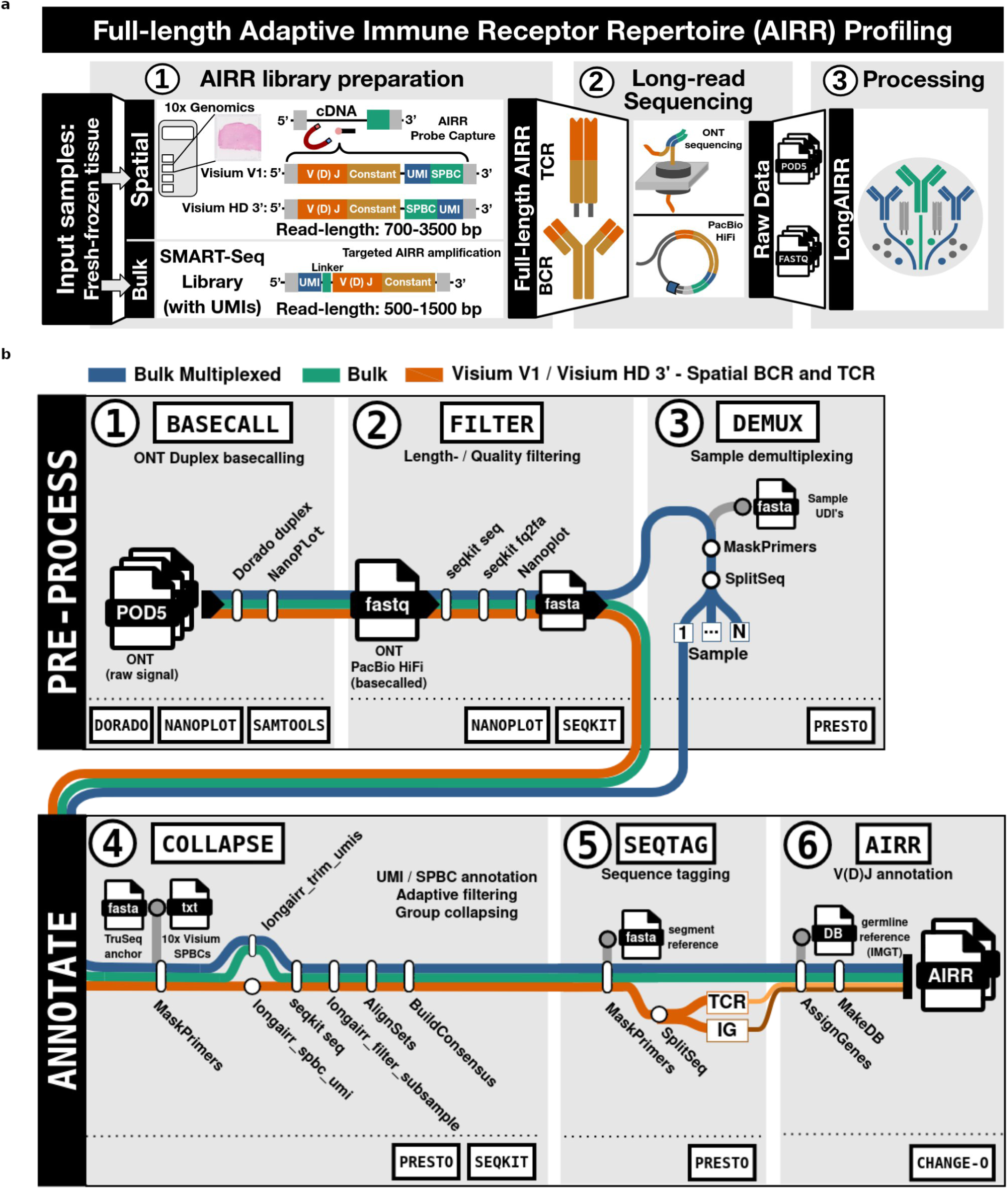
LongAIRR framework for experimental and computational profiling of adaptive immune receptor repertoires. **a:** Workflow for full-length adaptive immune receptor repertoire (AIRR) profiling using long-read sequencing technologies (Oxford Nanopore Technologies and Pacific Biosciences), applicable to spatial AIRR (Visium V1 and Visium HD 3’) or bulk AIRR analyses. **b:** Overview of *LongAIRR* software modules. Pre-processing modules (top): (1) *LongAIRR basecall*, ONT basecalling on raw data in ONT’s pod5 format, only for ONT data. (2) *LongAIRR filter,* length and quality filtering of FASTQ input files (from ONT or PacBio). (3) *LongAIRR demux* (optional), demultiplexing of multiplexed bulk samples. Annotation modules (bottom): (4) *LongAIRR collapse*, UMI and SPBC annotation, adaptive length filtering and consensus sequence generation from UMI-SPBC sequence groups. (5) *LongAIRR seqtag*, constant region annotation and splitting of BCR and TCR sequences into separated files. (6) *LongAIRR airr,* V(D)J annotation using a locus-specific reference database, producing AIRR-formatted TSV tables.

Because BCR and TCR transcripts can reach ∼3.5 kb in length (Lefranc2005), long-read sequencing is required to associate the receptor V(D)J region located at the 5′ end with the SPBC and UMI at the 3′ end of the read (**Fig. 1a**). Both ONT and PacBio long-read sequencing have been applied to full-length AIRR profiling (Benotmane2023, Engblom2023, Jiang2026, Luo2024). These sequencing technologies differ substantially in their sequencing chemistry and error profiles. ONT sequencing generates raw electrical signal data that is converted into nucleotide sequences through basecalling, whereas PacBio sequencing produces high-accuracy circular consensus reads (HiFi) by repeatedly sequencing the same molecule (Amarasinghe2020, Logsdon2020, Wenger2019). As a result, PacBio reads typically achieve higher per-read accuracy, while ONT accuracy depends on sequencing chemistry and basecalling algorithm (Amarasinghe2020, Hall2024, Logsdon2020). In addition to these technical differences, sequencing costs vary considerably depending on instrument access and consumable pricing, with PacBio sequencing typically representing the more expensive option. However, the extent to which full-length spatial AIRR transcriptomics data derived from these technologies are comparable remains largely unexplored.

Despite the increasing adoption of spAIRR sequencing, standardized computational workflows for analyzing long-read spatial AIRR datasets remain limited. Existing frameworks such as nf-core/airrflow (Gabernet2024) or the Immcantation framework (Gupta2015, VanderHeiden2014) address certain aspects of spAIRR analysis, but are not specifically designed for long-read spatial transcriptomics data. Consequently, most spAIRR studies to date have largely relied on study-specific or custom analysis workflows, which limits comparability between processed datasets and prevents systematic evaluation of computational parameters (Engblom2023, Jiang2026, Liu2022, Meylan2022). In addition, T cell receptor repertoire focused approaches such as SPTCR perform sequence reconstruction without incorporating UMI information (Benotmane2023), which restricts their suitability for B cell receptor analysis where somatic hypermutation must be distinguished from sequencing errors.

Here we present *LongAIRR*, a computational framework for the analysis of long-read spatial transcriptomics data, enabling robust AIRR annotation across sequencing platforms, including ONT and PacBio. *LongAIRR* is implemented as a modular command-line toolbox that integrates seamlessly with workflow managers such as Snakemake (Mölder2021) and provides interoperability with downstream AIRR analysis frameworks including Immcantation (Gupta2015, VanderHeiden2014). The framework supports both spatial and bulk AIRR datasets (**Fig. 1**).

We demonstrate that spAIRR data from simultaneous spatial profiling of BCR and TCR repertoires with either ONT R10 or PacBio sequencing is comparable in both quantity and quality. Across different experimental protocols, we further demonstrate that an adaptive filtering strategy, which dynamically refines read selection, enables accurate consensus reconstruction and mitigates platform-specific sequencing errors. This approach allows ONT-derived AIRR sequences to achieve accuracy comparable to PacBio HiFi reads in consensus sequences with at least three underlying raw reads. Finally, we derive evidence-based guidelines for the consistent and robust analysis of spatial AIRR data.

## RESULTS

In this study, we performed spAIRR sequencing on human glioma tissue samples using a capture-probe-based enrichment protocol targeting a comprehensive set of BCR and TCR transcripts (encoding Ig γ1–γ4, Ig α1-α2, Ig μ, Ig ε, Ig δ, Ig κ, Ig λ, TCR α, TCR β, TCR γ and TCR δ chains), which was developed based on previously described approaches (Benotmane2023, Engblom2023). spAIRR libraries were generated using 10x Genomics Visium V1 technology and the successor technology Visium HD 3′. To enable direct comparison of long-read sequencing technologies, libraries were sequenced using ONT MinION R10.4.1 (ONT R10) flow cells (S1_ONT, S2_ONT, S3_ONT, S4_ONT) or PacBio HiFi technology (S1_PB, technical replicate of S1_ONT, see **Table 1**). In addition, a previously published spatial TCR repertoire dataset generated using ONT R9.4 (ONT R9) chemistry (Benotmane2023), for which both raw and processed sequencing data are publicly available, was reprocessed with *LongAIRR*. Bulk AIRR sequencing was performed on another subset of samples to allow comparison across library preparation protocols.

**Table 1:** Overview of datasets, samples and experimental approaches.

| ID | Biological sample | Experiment type | Library preparation | Sequencing | Basecalling | Quality threshold |
| --- | --- | --- | --- | --- | --- | --- |
| S1_ONT_Q10 | tumor 1 | Visium V1 | BCR+TCR described in this study | ONT (R10) | Simplex + duplex | PHRED10 |
| S1_ONT_Q17 | tumor 1 | Visium V1 | BCR+TCR described in this study | ONT (R10) | Simplex + duplex | PHRED17 |
| S1_ONT_Q17_duplex | tumor 1 | Visium V1 | BCR+TCR described in this study | ONT (R10) | Duplex only | PHRED17 |
| S2_ONT_Q17 | tumor 2 | Visium V1 | BCR+TCR described in this study | ONT (R10) | Simplex + duplex | PHRED17 |
| S3_ONT_Q17 | tumor 3 | Visium V1 | BCR+TCR described in this study | ONT (R10) | Simplex + duplex | PHRED17 |
| S1_PB_Q17 | tumor 1 | Visium V1 | BCR+TCR described in this study | PacBio HiFi | / | PHRED17 |
| ONT_R9_longAIRR | PUB_01* | Visium V1 | Benotmane et al. | ONT (R9) | Benotmane et al. | PHRED17 |
| ONT_R9_original | PUB_01* | Visium V1 | Benotmane et al. | ONT (R9) | Benotmane et al. | Analysed data from publication |
| S4_ONT_Q17_HD | tumor 4 | Visium HD 3' | BCR+TCR described in this study | ONT (R10) | Simplex + duplex | PHRED17 |
| S5_ONT_Q17_UDI6 | tumor 5 | Bulk Takara | SMART-seq BCR (with UMIS) | ONT (R10) | Simplex + duplex | PHRED17 |
| S6_ONT_Q17_UDI20 | tumor 6 | Bulk Takara | SMART-seq BCR (with UMIS) | ONT (R10) | Simplex + duplex | PHRED17 |
| S7_ONT_Q17_UDI12 | tumor 7 | Bulk Takara | SMART-seq BCR (with UMIS) | ONT (R10) | Simplex + duplex | PHRED17 |
| S8_ONT_Q17_UDI13 | tumor 8 | Bulk Takara | SMART-seq BCR (with UMIS) | ONT (R10) | Simplex + duplex | PHRED17 |

\*Benotmane2023

### Sequence quality and yield in ONT and PacBio sequencing

The purified spAIRR library length-profile after capture probe-based enrichment and amplification ranged from 600 bp to more than 5,000 bp, with a peak around 1,400 bp (**Fig. 2a**). ONT and PacBio sequencing data was generated from the same input library (S1_ONT, S1_PB, see **Table 1**). The ONT R10 raw sequencing data generated in this study (**Fig. 2b, 2c**) showed a read length distribution comparable to previously published ONT R9 sequencing data for spatial TCR sequencing (**Fig. 2d**) (Benotmane2023). Read length distributions differed between sequencing platforms with PacBio producing fewer long reads (> 4000 bp) than ONT sequencing (**Fig. 2b-d**).

**Figure 2:**
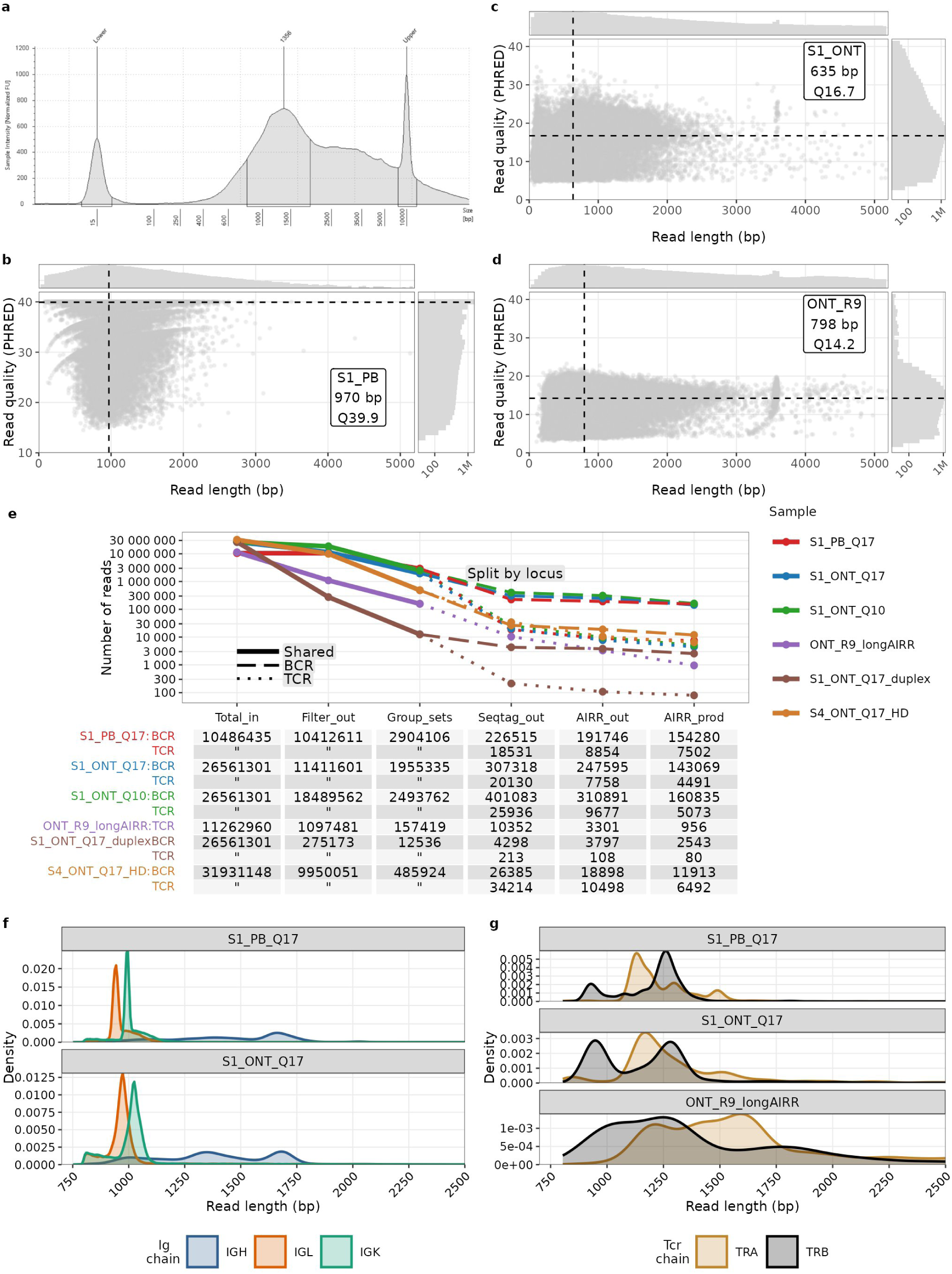
Comparable yield and sequence accuracy across ONT and PacBio datasets processed with LongAIRR. **a:** TapeStation profile of the probe-enriched and amplified spatial AIRR library (combined BCR and TCR) after post-capture amplification and before preparation of sequencing libraries. Peaks at 15 bp and 10,000 bp are not part of the library and correspond to the lower and upper markers**. b-d:** Sequencing quality (PHRED) and length distribution of raw sequence reads from (b) PacBio (S1_PB), (c) ONT R10 (S1_ONT) and (d) ONT R9 (ONT_R9). **e:** Number of sequences retained across *LongAIRR* processing steps: length- and quality-filtered reads (*Filter_out*), UMI-SPBC group consensus sequences (*Group_sets*), sequences with detected BCR or TCR constant sequence segments (*Seqtag_out*) and V(D)J-annotated AIRR sequences (all: *AIRR_out*; productive: *AIRR_prod*). Colors indicate datasets (see Table 1). Line types indicate loci: combined BCR and TCR reads (solid), BCR (large dashes) and TCR (small dashes) after constant region annotation and BCR/TCR splitting. **f**, **g:** Read-length distributions of productive receptor sequences (*AIRR_prod*) in BCR (f) and TCR (g) sequences. Headers in grey boxes indicate dataset identifier (see Table 1).

A major known difference between ONT R9, ONT R10, and PacBio sequencing is the raw read quality, commonly summarized by the PHRED score. PHRED scores represent the per-base error probabilities on a logarithmic scale, where each increase of 10 corresponds to a tenfold reduction in error rate, such that Q10 and Q20 correspond to estimated error rates of 10% and 1% respectively (Ewing1998). PacBio raw reads generated using circular consensus strategy showed the highest overall sequencing quality (median Q39.9, **Fig. 2b**). Basecalling of ONT R10 data was performed using the super-high-accuracy model (dna_r10.4.1_e8.2_400bps_sup@v5.0.0), which generates simplex and duplex reads (see Methods section *ONT Basecalling*). The resulting sequences reached higher quality values (median Q16.7, **Fig. 2c**) than previously published ONT R9 data processed with an earlier basecalling model (median Q14.2, **Fig. 2d**) (Benotmane2023).

To systematically compare ONT and PacBio sequencing and assess the effect of different quality filtering strategies on the yield and sequence quality of receptor sequences, we tracked sequence retention across *LongAIRR’s* processing modules and applied several quality thresholds (**Fig. 1b** and **Fig. 2e**). For ONT R10 data, we defined three quality thresholds: Q10 (simplex and duplex reads), Q17 (simplex and duplex reads), and Q17_duplex (duplex reads only). These filters retained 70% (18,489,562/26,561,301) of reads for S1_ONT_Q10, 43% (11,411,601/26,561,301) for S1_ONT_Q17, and 1% (275,173/26,561,301) for S1_ONT_Q17_duplex, calculated from *Total_in* to *Filter_out* (**Fig. 2e**).

### Consensus sequence building and annotation of AIRR sequences

Following initial filtering, *LongAIRR* annotates UMI and SPBC in each read. Reads lacking valid SPBC sequences are removed, while the remaining reads are grouped based on UMI-SPBC combinations, corresponding to amplifications of the same originally captured mRNA molecule (**Fig. 1b**, **Module 4**). Reads within each UMI-SPBC group are aligned using multiple sequence alignment (MSA) and subsequently collapsed into one consensus sequence to mitigate sequencing errors. The resulting number of distinct groups for each dataset and quality filtering strategy is reported in the *Group_sets* column (**Fig. 2e**). Up to this stage, using the *LongAIRR* modules to process long-read data is not restricted to AIRR transcripts and can be applied to other long-read transcriptomics sequencing datasets containing comparable read structures.

Next, consensus sequences originating from BCR or TCR transcripts were identified by mapping short constant region reference sequences to the consensus sequences (**Fig. 1b**). At this stage, sequences are separated by receptor locus (BCR or TCR), as visualized in the locus-specific dashed lines in Fig. 2e. The proportion of consensus sequences containing identifiable BCR or TCR constant regions was comparatively low (*Seqtag_out*, *Group_sets*, **Fig. 2e**): 8.4% for S1_PB_Q17 (PacBio) (225,237 BCR sequences and 18,515 TCR sequences for a total of 2,904,106 consensus sequences) and 6.5% for the reprocessed public ONT_R9_longAIRR TCR repertoire dataset (10,352 TCR sequences for 157,419 consensus sequences). For ONT R10 data, the proportion of BCR and TCR sequences per total consensus sequences increased with filtering stringency: 17.1% for S1_ONT_Q10 ((401,083 BCR + 25,936 TCR)/2,493,762 Group sets), 16.7% for S1_ONT_Q17 ((307,318 + 20,130)/1,955,335), 35.9% for S1_ONT_Q17_duplex ((4,298+213)/12,536). Similar retention statistics were observed across additional spatial Visium V1 (**Fig. S1a**) and Visium HD 3′ samples (S4_ONT_Q17_HD, **Fig. 2e**). In contrast, AIRR libraries generated using bulk level PCR amplification with specific primers of BCR-transcripts show substantially higher receptor recovery rates, with 43-94% of UMI groups containing identifiable receptor constant regions (**Fig. S1b**). These results indicate that the spAIRR library after probe-based enrichment still contains a substantial amount of transcripts other than BCR or TCR transcripts.

Consensus sequences that contain adaptive immune receptor constant regions (*Seqtag_out*, **Fig. 2e**) were next processed with *IgBLAST* (VanderHeiden2014, Ye2013) to assign V, D and J gene segments and reconstruct receptor rearrangements (*AIRR_out*, **Fig. 2e**, **Fig 1b**). This step generates AIRR-formatted receptor annotations and includes non-productive sequences or such with incomplete AIRR annotations. The proportion of productive sequences (*AIRR_prod*, *AIRR_out*, **Fig. 2e**) was highest for S1_PB_Q17 (PacBio): 80.4% BCR (154,280 reads in AIRR_prod / 191,746 reads in AIRR_out) and 84.7% TCR (7,502/8,854). For ONT R10 data, S1_ONT_Q17_duplex achieved 66.9% BCR (2,543/3,797) and 74.0% TCR (80/108), while S1_ONT_Q17 simplex-duplex yielded 57.7% BCR (143,069/247,595) and 57.8% TCR (4,491/7,758). These productivity rates are consistent with the expected sequencing accuracy of the platforms, as higher sequencing accuracy in PacBio sequencing reduces the number of frame-shift or missense errors and increases the fraction of in-frame receptor sequences (Hall2024, Logsdon2020, Wenger2019). The length distributions of these productive sequences (*AIRR_prod*, **Fig. 2e**) are similar between sequencing platforms and match the expected transcript length for BCR and TCR sequences (**Fig. 2f-g**) (Lefranc2005).

Together, these results illustrate the pronounced trade-off between sequencing yield and read quality in ONT R10 data. In summary, the Q17 quality filtering approach including simplex and duplex reads provided a good balance between consensus sequence yield and sequence quality. In following analyses, we therefore focus on comparing PacBio and ONT data using the filtering settings S1_PB_Q17, S1_ONT_Q10 and S1_ONT_Q17.

Across all datasets, BCR sequences were more abundant than TCR sequences, consistent with previous reports (Engblom2023). This likely reflects the higher number of immunoglobulin transcripts in antibody-secreting plasma cells and B cells compared to the lower abundance of TCR transcripts in T cells.

### Accuracy of UMI and SPBC annotation

The number of remaining sequencing reads after initial length and quality filtering typically exceeds the number of reconstructed UMI-SPBC groups by more than threefold (*Filter_out*, *Group_sets*, **Fig. 2e**), indicating that some groups are represented by multiple reads.

It has previously been reported that UMI assignment can be ambiguous and that there may be a bias towards poly-T UMIs in Visium V1 data (Engblom2023). To assess the accuracy of UMI and SPBC annotation, we examined cases where a single UMI sequence was associated with multiple distinct SPBCs. Such events are theoretically highly unlikely given the diversity of a 12 nt UMI sequence (4^12^ = 16.7 million possible combinations). In the case of Visium V1 technology in combination with ONT R10 sequencing (S1_ONT_Q17), fewer than 2% of observed UMI-SPBC pairs were associated with multi-SPBC UMI groups across all loci. In contrast, Visium V1 technology in combination with PacBio (S1_PB_Q17) shows slightly higher rates, exceeding 2% for IGK and IGL loci (**Fig. S2a**). UMI sequences shared between multiple SPBC were predominantly composed of poly-T sequences, further suggesting sequencing artifacts or chimeric molecule formation (**Fig. S2b**).

For Visium HD 3′, the number of ambiguous UMI-SPBC groups was higher (**Fig. S2a**), but there was no enrichment for single ambiguous UMIs and no observable sequence bias in the UMIs with multiple SPBCs (**Fig. S2b**). This suggests that due to the higher number of SPBCs (4992 in Visium V1, >11 million in Visium HD 3′) and due to a different experimental design of the capture primers, same UMIs in combination with different SPBCs in Visium HD 3′ data represent distinct captured mRNA molecules rather than technical artifacts.

Based on these results, we removed UMI-SPBC groups containing poly-T UMI sequences, as reported by Engblom et al (Engblom2023). In addition, sequences showing ambiguous locus assignments or uncertain nucleotide-annotations (“N” in the consensus building) in the variable region, were excluded from subsequent analyses.

### Adequate sequencing coverage across UMI-SPBC groups

With these filtering criteria applied, we characterized the distribution of reads per unique UMI-SPBC group in both ONT and PacBio processed datasets (**Fig. 3a**). Group size was determined using the *N_ORIG* column of the spatial AIRR output generated by *LongAIRR*, which corresponds to the number of reads assigned to the same UMI-SPBC group after the pre-processing modules (**Fig. 1b**, **Module 4**).

**Figure 3:**
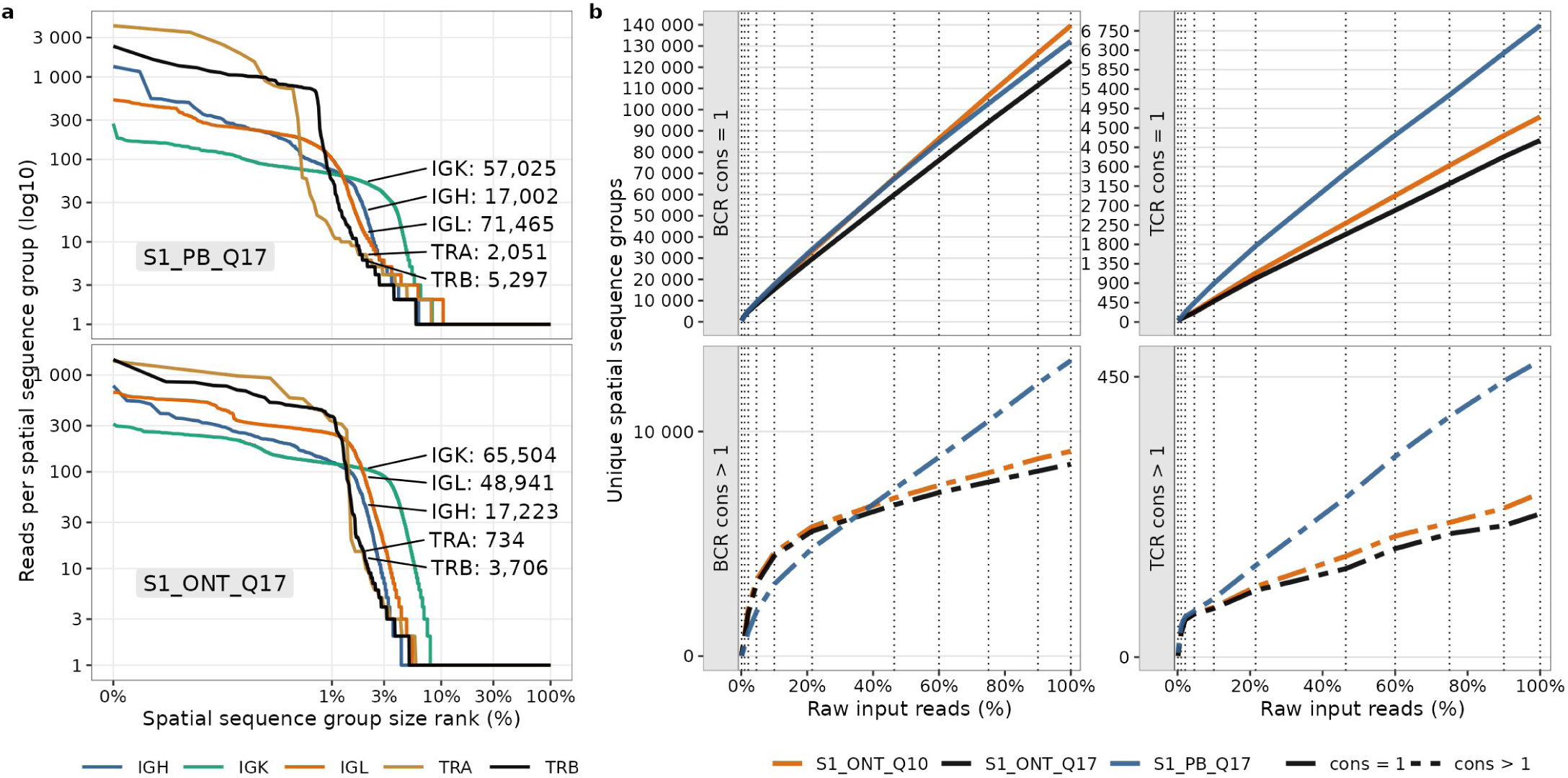
Sequencing coverage of UMI-SPBC groups at different sequencing depths. **a:** Number of reads per UMI-SPBC consensus group in S1_PB_Q17 (top) and S1_ONT_Q17 (bottom) datasets. Consensus groups are ranked by size and the ranks are displayed on a percentile scale (x-axis). Numbers indicate the total number of consensus groups per transcript type (encoding Ig heavy, Ig κ, Ig λ, TCR α, TCR β chains). **b:** Number of productive receptor sequences for BCR (left) and TCR (right) identified in subsampled datasets. Datasets are subsampled step-wise from 100% to 0.1% of raw input sequences. UMI-SPBC are split according to the number of supporting reads: single-read UMI-SPBC groups (cons = 1, top row, solid); UMI-SPBC groups supported by > 1 read (cons > 1, bottom row, dash-dotted). Colors indicate datasets (see Table 1).

The number of sequences per UMI-SPBC group followed similar distributions across sequencing technologies and transcript family (Ig heavy chain, Ig κ, Ig λ, TCR α, TCR β). The largest TCR α UMI-SPBC groups made up approximately 1% of all unique TCR α UMI-SPBC groups and contained more reads in the PacBio (S1_PB_Q17) dataset (∼3,000 reads) compared to the ONT dataset (S1_ONT_Q17) (∼1,000 reads) (**Fig. 3a**). Between 2-8% of the UMI-SPBC groups contained between 3 and 100 reads. The majority (∼90%) of groups contained only a single read (**Fig. 3a**). spAIRR data generated with the Visium HD 3′ platform is comparable, with large group sizes (>1,000 reads) across all loci and the majority of groups with a size of 1 (∼90%) (S4_ONT_Q17_HD in **Fig S3d**).

To assess whether a higher sequencing depth would increase the number of UMI-SPBC groups supported by more than one read, we performed subsampling analysis on the input fasta sequences and processed each subset of fasta files with *LongAIRR* (**Fig. 3b**). While the total number of UMI-SPBC combinations with only one read increased linearly with sequencing depth, the number of UMI-SPBC groups with at least 2 reads did not substantially increase with sequencing depth (S1_ONT_Q10, S1_ONT_Q17, S1_PB_Q17) (**Fig. 3b**). Consequently, higher sequencing depth only marginally increases the number of UMI-SPBC groups supported by more than one read. These findings also indicate that the majority of UMI-SPBC groups with only one read likely represent technical artifacts rather than true mRNA molecules.

### Sequence accuracy in larger UMI-SPBC groups is comparable between PacBio and ONT R10 sequencing

*LongAIRR* implements consensus building per UMI-SPBC group to improve overall sequence accuracy and mitigate potential sequencing errors that are randomly distributed across the sequence. A key challenge in consensus building is that reads within the same group may vary substantially in read-length due to incomplete DNA amplification, premature sequencing termination, sequencing read-through or contamination by unrelated read artifacts. MSA of reads with very different lengths introduced gaps and unexpected insertions/deletions negatively impacting the consensus sequence accuracy, specifically for UMI-SPBC groups with sometimes thousands of reads.

To reduce the read-length variation within UMI-SPBC groups, we developed an adaptive length filtering approach that identifies the most abundant read length and retains only reads within a 5% margin around this length (**Fig. 4a, 4b, 1b Module 4**). Adaptive length filtering is applied to every UMI-SPBC group containing more than four reads and leads to an increase of productive AIRR sequences by more than 10% across multiple datasets (**Fig. 4c**).

**Figure 4:**
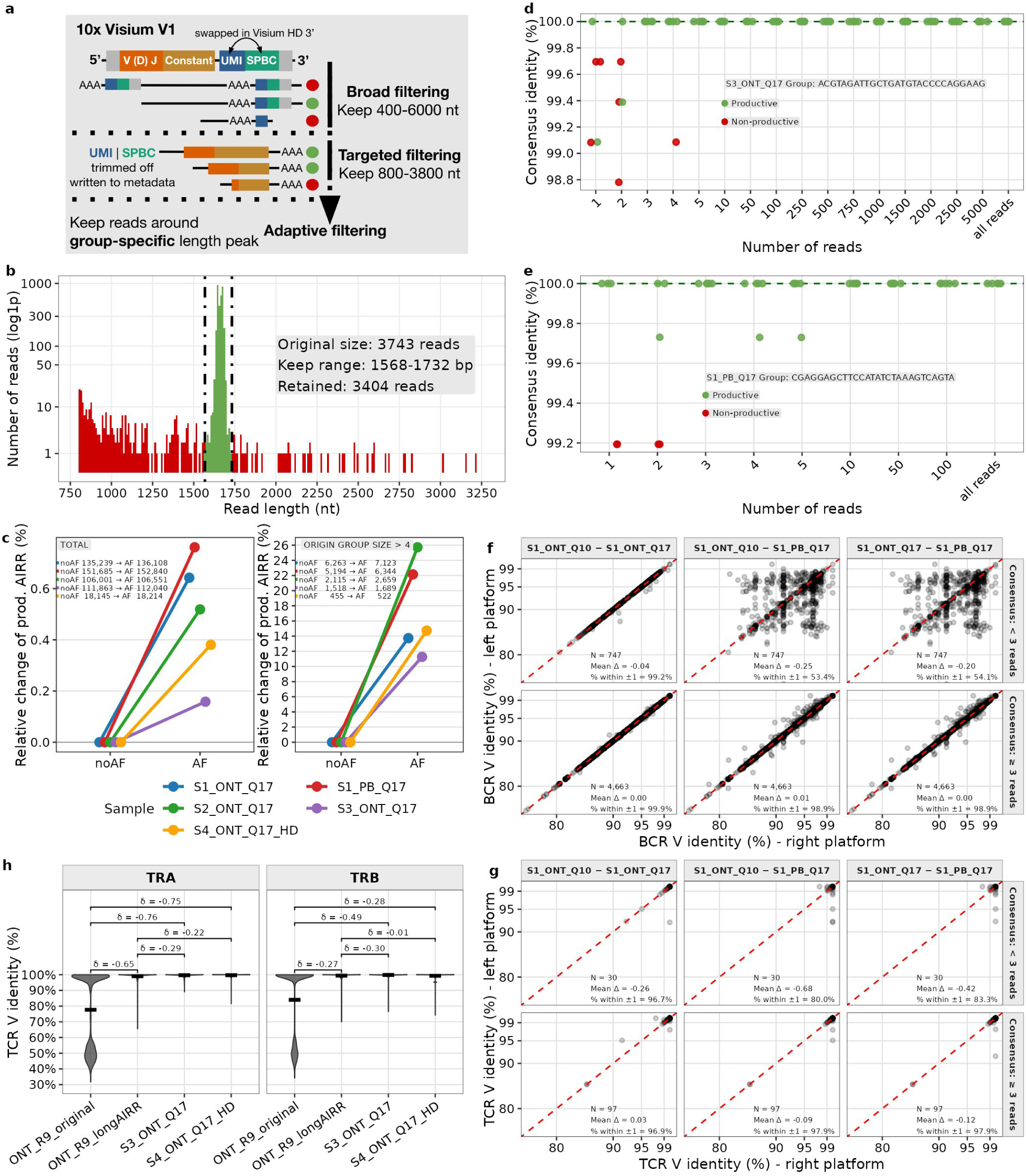
*LongAIRR* uses adaptive filtering to improve consensus building. **a:** Schematic of the read filtering strategy in *LongAIRR*, prior to consensus building. Fixed-length filtering (consisting of broad filtering before and targeted filtering after UMI-SPBC assignment and trimming) is followed by adaptive filtering only retaining reads around the most abundant read length peak per consensus group. Exemplary read structure shown for Visium V1 reads. **b:** Read length distribution of a representative UMI-SPBC group (dataset: S3_ONT_Q17). Dotted lines indicate 5% margin around the detected peak used for adaptive filtering. Reads within the window are retained (green), while outliers are discarded (red). Read counts are shown on a log1p scale (y-axis). **c:** Relative change of number of productive sequences after adaptive filtering (AF), compared to no adaptive filtering (noAF). Changes are shown for all groups (left) and groups with > 4 reads prior to filtering (right). Numbers in each figure indicate the absolute increase of productive sequences. **d**, **e:** Comparison of full-group consensus to consensus sequences built on subsamples of UMI-SPBC read groups. A representative UMI-SPBC group from (d) S3_ONT_Q17 and (e) S1_PB_Q17 dataset is shown. Sequence identity is calculated as percentage identify using the hamming distance within the FWR1-FWR4 region. UMI- SPBC group identifiers are indicated in the panels. Colors indicate productivity status of the consensus receptor sequence: productive (green); non-productive (red). **f**, **g:** Pairwise comparison of V-region identity for UMI-SPBC consensus groups matched across datasets (S1_PB_Q17, S1_ONT_Q17, S1_ONT_Q10). Comparisons are shown separately for BCR (f) and TCR (g). Each point represents one matching UMI-SPBC group, only productive consensus sequences from UMI-SPBC groups present in all 3 datasets are shown. y-axis and x-axis indicate the V identity in the first and second dataset indicated in the column title respectively (horizontal grey box). Groups are stratified by consensus group size: < 3 reads in at least one dataset (top); ≥ 3 reads in all three datasets (bottom). N: number of shared groups, Δ: mean difference between V identities. The proportion of groups differing by no more than 1% in V-region identity is indicated. **h:** TCR V-region identity across datasets for TRA (left) and TRB (right). Datasets: ONT_R9_original (ONT R9; Visium V1; SPTCR processing), ONT_R9_longAIRR (ONT R9; Visium V1; *LongAIRR* processing), S3_ONT_Q17 (ONT R10; Visium V1; *LongAIRR* processing) and S4_ONT_Q17_HD (ONT R10; Visium HD 3’; *LongAIRR* processing). Pairwise differences between datasets are quantified using Cliff’s delta (δ).

### Consensus building improves sequence accuracy independent of sequencing technology

In a next step, we aimed to evaluate the sequence accuracy in single-read UMI-SPBC groups and test the robustness of consensus building for small UMI-SPBC groups. To this end, we examined how downsampling of UMI-SPBC groups affects the consensus sequence (**Fig. 4d, 4e**, **S3a**, **S3b**). Five independent subsamples with sizes between 1 read and 5,000 reads were derived from representative large UMI-SPBC groups in the S3_ONT_Q17 (**Fig. 4d**) and PacBio datasets (**Fig. 4e**). The consensus sequence obtained from each subsample was compared to the consensus sequence generated from the full UMI-SPBC group by calculating the percentage of sequence identity. Sequence similarity was calculated only for the variable receptor region spanning FWR1–FWR4, while flanking adapter and constant regions were excluded from the analysis. Of note, *LongAIRR* applies a similar subsampling strategy that limits large UMI–SPBC groups to a maximum of 750 reads per consensus to reduce computational cost without compromising consensus sequence quality (**Fig. S3c**, **Table S2**).

Subsamples with one or two reads showed the lowest mean sequence identity of 99.48% in the ONT-R10-derived group and 99.65% in the PacBio-derived group (**Fig. 4d**, **4e**). This behavior was also consistently observed in other subsampled UMI-SPBC groups (**Fig 4d, 4e**, **Fig. S3a**, **S3b**). This result suggest that accurate consensus sequence building requires at least 3 sequences, independent of the sequence quality achieved. This is likely due to the presence of PCR amplification errors in the raw sequencing reads, that can also be corrected by consensus building.

This conclusion is further supported when comparing the V-region identity of productive receptor sequences generated by *LongAIRR* based on PacBio and ONT data (**Fig. 4f, 4g**). V-region identity was determined during V(D)J annotation (**Fig. 1b**, **Module 6**). We matched UMI-SPBC consensus sequences across three datasets (S1_ONT_Q10, S1_ONT_Q17, S1_PB_Q17) and compared the V-region identities for BCR and TCR sequences (**Fig. 4f, 4g**).

Consensus sequences supported by fewer than three reads showed poor agreement between ONT and PacBio datasets (**Fig. 4f, 4g**, **top rows**). Only 54.1% of BCR consensus sequences from matching UMI-SPBC groups (S1_ONT_Q17, S1_PB_Q17) and 83.3% of TCR consensus sequence pairs (S1_ONT_Q17, S1_PB_Q17) showed a difference in V-region identity of less than 1%. V-region identity was more similar between consensus sequences supported by at least three reads across all dataset pairings (**Fig. 4f, 4g**, **bottom rows**). At least 98.9% of matched productive BCR consensus sequence pairs differed by less than 1% in V-region identity, compared to 97.9% for TCR consensus sequences. In this analysis, S1_ONT_Q17 and S1_ONT_Q10 show very similar results, since the underlying raw data is the same only sequences overlapping between both datasets were included.

### *LongAIRR* achieves higher sequence accuracy than previous approaches

To further validate the sequence accuracy achieved by *LongAIRR* and compare our methodology to existing analysis approaches, we reanalyzed a previously published TCR spatial sequencing dataset generated using ONT R9 chemistry (ONT_R9_original), which was processed using the *SPTCR* pipeline in the original study (Benotmane2023). Reprocessing the same dataset with *LongAIRR* substantially increased the observed V-region identity for both TCR α and TCR β transcripts compared to the published *SPTCR* output, as indicated by Cliff’s delta (δ) values of −0.65 and −0.27 for TCR α and TCR β transcripts respectively (**Fig. 4h**, Methods) (Cliff1993). TCR sequences generated in our study using ONT R10 sequencing and processed with *LongAIRR* (S3_ONT_Q17 and S4_ONT_Q17_HD) showed the highest overall V-region identity distributions across both loci (**Fig. 4h**).

These results demonstrate that *LongAIRR* substantially improves the accuracy of AIRR sequence reconstruction from ONT data and yields V-region identity distributions comparable across sequencing chemistries. Raw sequencing reads are affected by both platform-specific errors and PCR amplification artifacts. However, consensus sequences supported by at least three reads achieved sufficient accuracy, largely independent of the sequencing technology.

### Recommendations for consensus count filtering for spatial AIRR analysis

While consensus-based error correction achieves good sequence accuracy for consensus counts > 2 in both ONT and PacBio datasets (**Fig. 4**), spatial localization of sequences represents an additional analytical challenge. spAIRR data exhibits inherent sparsity, with individual clones captured in only a small fraction of SPBCs. This biological characteristic is further complicated by a technical challenge: 90% of consensus sequences have consensus count = 1 (**Fig. 3a**, **S3d**), suggesting that a substantial proportion of sequences where UMI and SPBC associations likely result from sequencing errors or library preparation artifacts rather than true spatial transcript barcoding. To determine the optimal balance between removal of these technical artifacts and preservation of biological signal in downstream spatial analysis, we systematically evaluated four filtering approaches and applied them to a Visium V1 (S1_ONT_Q17) and a Visium HD 3′ (S4_ONT_Q17_HD) dataset (**Fig. 5a**).

**Figure 5:**
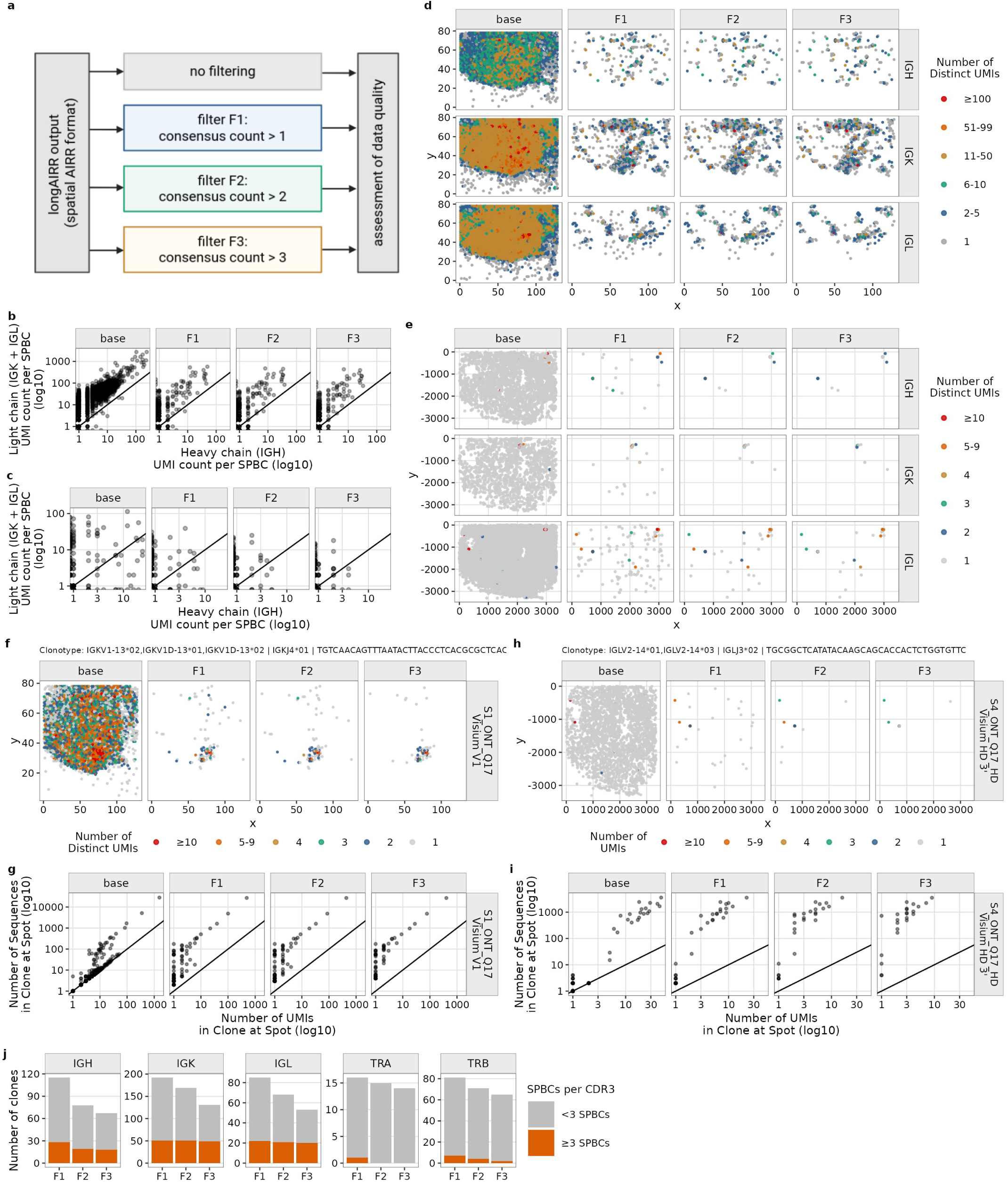
Trade-off between background removal and information retention in spatial AIRR data. **a:** Filtering approaches for spatial AIRR data. F1 removes UMI-SPBC consensus groups supported by a single read; F2 removes groups with ≤ 2 supporting reads; F3 removes groups with ≤ 3 supporting reads. **b**, **c:** Correlation between the number of UMIs per SPBC for light chain sequences (combined IGK and IGL; y-axis) and heavy chain sequences (IGH; x-axis). Each point represents one SPBC. Data are shown for (b) S1_ONT_Q17 (Visium V1) and (c) S4_ONT_Q17_HD (Visium HD 3’). **d**, **e:** Number of UMIs captured per Visium spot. Each point corresponds to one spot at its spatial coordinates. Colors indicate the number of UMIs per SPBC. Panels are faceted by transcript type (Ig heavy, Ig κ, Ig λ, rows) and filtering strategy (base, F1-F3; columns). Data are shown for (d) S1_ONT_Q17 (Visium V1) and (e) S4_ONT_Q17_HD (Visium HD 3’). **f**, **h:** Number of UMIs captured per Visium spots for two representative Ig light chain clones in (f) S1_ONT_Q17 (Visium V1) and (h) S4_ONT_Q17_HD (Visium HD 3’). Each point corresponds to one spot at its spatial coordinates. Colors indicate the number of UMIs per SPBC. Panels are faceted by filtering strategy (base, F1-F3). Clones are defined by identical V-J annotation and CDR3 nucleotide sequence with the clone identifiers indicated in the panel subtitles. **g**, **i:** Correlation between the number of UMIs per SPBC (x-axis) and the total number of reads contributing to consensus sequences per SPBC (y-axis) for the same representative Ig light chain clones as in (f, h). Each point represents one SPBC. Panels are faceted by filtering strategy (base, F1-F3). Data are shown for (g) S1_ONT_Q17 (Visium V1) and (i) S4_ONT_Q17_HD (Visium HD 3’). **j:** Number of unique CDR3 nucleotide sequences retained after filtering (F1-F3), faceted by transcript type (Ig heavy, Ig κ, Ig λ, TCR α, TCR β). Colors indicate whether CDR3 sequences are detected in ≥ 3 SPBCs.

The four filtering approaches differ by the minimum number of sequences per consensus: ‘Base’ (unfiltered), F1 (consensus count > 1), F2 (consensus count > 2), and F3 (consensus count > 3). F2 is the filtering strategy that is required for optimal sequence quality (**Fig. 4**). F1 represents a less stringent approaches that may better preserve spatial information in sparse datasets. The first filtering criterion (F1) reduced the number of unique UMI-SPBC combinations and the number of SPBC with annotated spAIRR sequences substantially (30 fold for the number of IGH consensus sequences, **Table 2**), in line with the hypothesis that sequences with consensus count of 1 predominantly represent technical artifacts rather than legitimate biological signal.

**Table 2:**
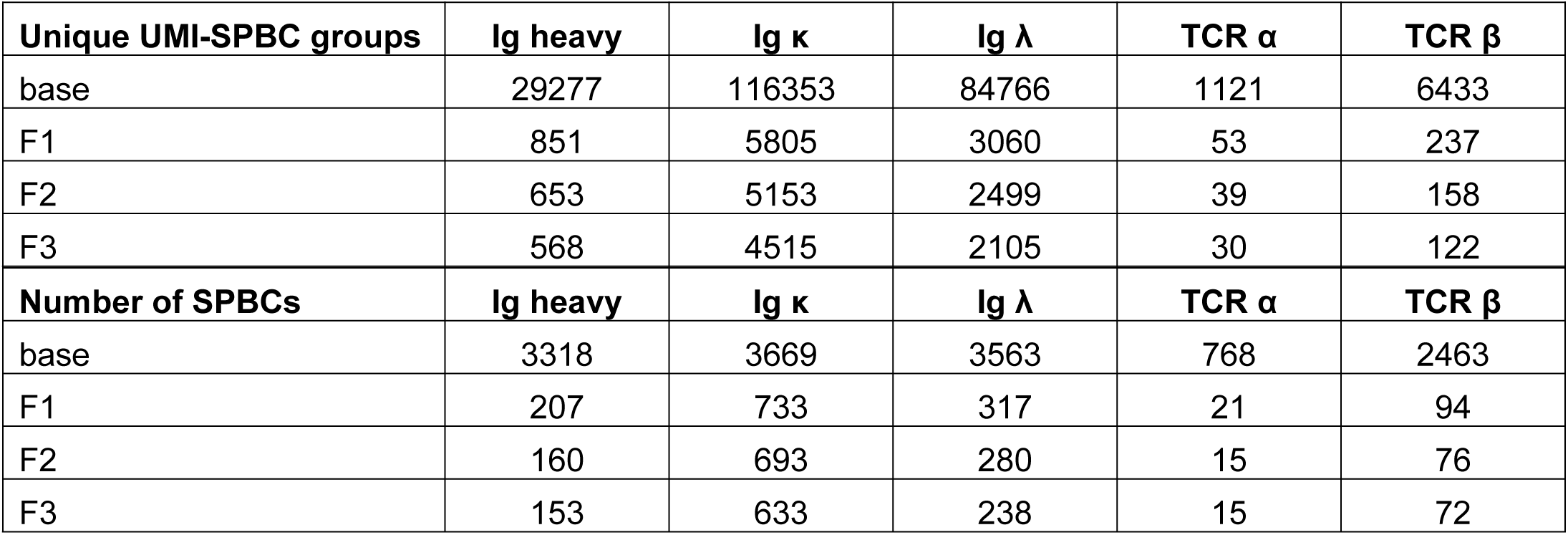
Number of unique UMI-SPBC groups and SPBCs with spAIRR sequences after different filtering approaches. The table displays data for the S1_ONT_Q17 dataset only (Visium V1).

| Unique UMI-SPBC groups | Ig heavy | Ig $\kappa$ | Ig $\lambda$ | TCR $\alpha$ | TCR $\beta$ |
| --- | --- | --- | --- | --- | --- |
| base | 29277 | 116353 | 84766 | 1121 | 6433 |
| F1 | 851 | 5805 | 3060 | 53 | 237 |
| F2 | 653 | 5153 | 2499 | 39 | 158 |
| F3 | 568 | 4515 | 2105 | 30 | 122 |
| Number of SPBCs | Ig heavy | Ig $\kappa$ | Ig $\lambda$ | TCR $\alpha$ | TCR $\beta$ |
| base | 3318 | 3669 | 3563 | 768 | 2463 |
| F1 | 207 | 733 | 317 | 21 | 94 |
| F2 | 160 | 693 | 280 | 15 | 76 |
| F3 | 153 | 633 | 238 | 15 | 72 |

The Visium V1 dataset showed a higher number of unique UMIs per SPBC than the Visium HD 3′ dataset, which is mainly due to the much higher resolution and therefore smaller number of cells per SPBC in the Visium HD 3′ dataset (**Fig. 5b, 5c**). Consequently, the heavy and light chain expression per SPBC showed robust positive correlation across all filtering conditions F1, F2 and F3 in the Visium V1 dataset, but was less pronounced in the sparser Visium HD 3′ dataset (**Fig. 5b, 5c**). Nevertheless, the spatial correlation of heavy and light chain sequences and their enrichment in confined tissue areas is apparent in both datasets, in line with the presence of locally defined tissue areas with higher immune cell infiltration (**Fig. 5d, 5e**). For Visium V1, we used gene expression profiles to locate immune cell aggregates, also known as tertiary lymphoid structures (TLS) and previously identified in gliomas (Cakmak2025), and confirmed that the percentage of TLS spots with available spAIRR data was comparable across approaches F1, F2 and F3 (**Fig. S4a**).

To further test the filtering approaches, we examined the spatial distribution of two representative clones across all four filtering conditions (**Fig. 5f-i**). In the unfiltered base condition, sequences of the clones where found across many SPBCs across the tissue section. In many of these SPBCs, the number of UMIs was identical to the number of sequences in a spot (UMI-to-sequence ratio = 1.0), a pattern characteristic of random spatial pattern rather than genuine clonal localization (**Fig. 5g, 5i**). Filtering condition F1 (consensus count > 1) effectively eliminated the randomly distributed background signal observed in the base condition, yielding spatially contiguous clone distributions consistent with biological expectations. Applying F2 and F3 removed some potentially legitimate sequences from SPBCs with lower UMI capture. Importantly, the additional sequence locations found in F1 did not substantially affect the distribution of mutual information between chains compared to F2 and F3 (**Fig. S4b**). Filtering approach F1 recovered the highest number of unique CDR3 sequences and the greatest number of CDR3 clonotypes spanning at least 3 SPBCs (**Fig. 5j**). The absolute number of UMI-SPBC groups and the number of SPBCs with annotated spAIRR data decreased progressively from F1 through F3 (**Table 2** and **Fig. 5d, 5e**).

In summary we conclude, that using F1 instead of F2 or F3 filtering approaches allow to recover more valid clonal spatial information and preserve biologically meaningful spatial patterns in sparse datasets. While F2 should be applied to obtain optimal sequence quality, especially for assessment of somatic hypermutations, F1 may represent a better starting point for any downstream analyses employing spatial statistics.

## DISCUSSION

spAIRR sequencing using a methodological combination of Visium-based spatial barcoding and long-read sequencing enables spatial mapping of B cell and T cell receptor repertoires within tissue sections (Benotmane2023, Engblom2023, Guo2025, Hudson2022, Liu2022, Ly2025, Meylan2022, Sudmeier2022). In this study, we use for the first time ONT R10 sequencing for simultaneous sequencing of spatially barcoded BCR and TCR sequences and we demonstrate that the resulting spAIRR sequences are comparable in quantity and quality to results obtained using PacBio sequencing.

The generation of high-quality spAIRR data for long-read sequencing output is enabled by *LongAIRR*, a new computational method that we introduce in this study. *LongAIRR* enables robust and reproducible processing across different library preparation protocols (Bulk, Visium V1 and the new Visium HD 3′) and sequencing platforms (ONT R9, ONT R10, PacBio HiFi). Despite substantial differences in input read quality and sequencing characteristics (Amarasinghe2020, Logsdon2020, Hall2024, Wenger2019), processing with *LongAIRR* yields consistent retention rates and comparable numbers of productive receptor sequences across different datasets. UMI-SPBC group size distributions were similar between the replicated ONT and PacBio dataset (S1_ONT_Q17, S1_PB_Q17), with the majority of UMI-SPBC groups supported by a single read and a smaller fraction supported by multiple reads (**Fig. 3**).

Rarefaction analysis indicated that all relevant receptor sequences are comprehensively covered in the small fraction of consensus sequences with more than one read and that increasing sequencing depth would not increase the number of total valid receptor sequences (**Fig. 3**). The sparsity of the resulting data therefore seems to be due to limits in capture efficiency in the Visium technology. From an experimental point of view, improving the probe-based enrichment of AIRR sequences in the sequencing libraries would allow a reduction in the number of total sequencing reads needed to reproduce similar spAIRR coverage.

The sequence rarefaction analyses and spatial analyses (**Fig. 3** and **5**), suggest that the majority of reads with a consensus support of n = 1 are technical artifacts, that likely reflect amplification chimeras or sequencing errors. We therefore recommend considering only consensus sequences with at least two supporting reads for spatial analyses such as spatial statistics and inference of spatial clone patterns.

In addition to the spatial information content, sequence quality is another analytical challenge that becomes especially important when looking for reliable somatic hypermutation data in BCRs. In this study we show that the accuracy of receptor sequences not only depends on raw sequencing read quality, but is also likely influenced by PCR-amplification errors. Consensus building is thus key to reliable spAIRR sequence quality (**Fig. 4**). We recommend only using consensus sequences with at least three supporting reads for critical analyses such as mutation calling or clonal lineage building. When applying these thresholds, the differences between PacBio and ONT based sequencing become negligible.

Our results position *LongAIRR* as a standardized framework for long-read spatially-resolved AIRR data analysis that enables consistent interpretation across sequencing platforms and experimental designs. By combining UMI- and SPBC-aware sequence grouping with adaptive filtering and consensus-based reconstruction, *LongAIRR* enables accurate recovery of combined BCR and TCR repertoires from ONT data at a level comparable to PacBio sequencing. The modular software architecture is a foundation for spatial AIRR profiling across current and emerging experimental techniques and existing downstream analysis workflows (Gupta2015, Hoehn2022, Nouri2018, Nouri2020, Stern2014, VanderHeiden2014) and position *LongAIRR* as a generalizable framework for cross-platform spatial immunogenomics. While developed for spatial AIRR profiling, core components of *LongAIRR* are applicable to other long-read sequencing workflows requiring UMI-based consensus reconstruction.

## MATERIAL AND METHODS

### EXPERIMENTAL METHODS

#### Ethical statement

Clinical data and tumor samples were obtained upon informed consent of the patients. The study was conducted in accordance with ethical standards such as the Declaration of Helsinki. The study protocol was endorsed by the ethical committee of the University Cancer Center Frankfurt (SNO-4-2022 and SNO-3-2025).

#### Sample collection and preparation

Resected adult-type diffuse IDH-wildtype and IDH-mutant glioma tissue samples were immediately stored at 4°C and transported on ice to the laboratory within 1–3 hours after surgical resection. The resected tumor tissue was embedded in OCT Tissue Tek, snap-frozen in liquid nitrogen, and stored at −80°C until further processing.

In addition, fresh frozen tissue from adult-type diffuse IDH-wildtype and IDH-mutant gliomas, which had been previously classified as tertiary lymphoid structure positive (Cakmak2025), were provided by the UCT BioBank. The fresh frozen samples were embedded in OCT Tissue Tek, and stored at −80°C until further processing.

#### 10x Visium library preparation and Illumina sequencing

Fresh-frozen glioma tumor sections were sectioned at 10 µm thickness and mounted onto slides from the Visium Spatial Gene Expression Slide & Reagent Kit (10x Genomics, USA). Tissue images were acquired at 20× magnification using the Vectra Polaris slide scanner (Akoya Biosciences, USA). Spatially barcoded cDNA synthesis and gene expression (GEX) library preparation were performed according to the manufacturer’s protocol (CG000239, 10x Genomics). Final GEX libraries were sequenced on a NextSeq1000/2000 sequencer (Illumina, USA) using P2, P3, or P4 reagent kits (100 cycles, XLEAP-SBS chemistry; Illumina, United States).

#### 10x Visium HD 3‘ library preparation and Illumina sequencing

Fresh-frozen glioma tumor sections were cut at 10 µm thickness and mounted onto glass slides (Superfrost Plus, ThermoFisher Scientific, Germany) for H&E staining according to manufactureŕs protocol (CG000804, 10x Genomics, USA) and tissue images were acquired at 20× magnification using the Vectra Polaris slide scanner (Akoya Biosciences, USA). Spatially barcoded cDNA synthesis and GEX library preparation were performed according to the manufactureŕs protocol (CG000805, 10x Genomics). Final GEX libraries were sequenced on a NextSeq 1000/2000 sequencer (Illumina, USA) using P4 reagent kits (100 cycles, XLEAP-SBS chemistry; Illumina, United States).

#### Spatial AIRR library preparation and ONT sequencing

For spatial AIRR sequencing, full length spatially barcoded cDNA generated within the 10x Visium Spatial Gene Expression workflow was used, similar to the methodologies described by Benotmane et al 2023 (Benotmane2023), and Engblom et al 2023 (Engblom2023).

##### Pre-capture amplification

Up to 100 ng of cDNA in 20 µl nuclease free water (NfW) was amplified prior to target enrichment using 25 µl of KAPA HiFi HotStart Ready Mix (2X) (Roche), and 7.5 µl of 10x cDNA primer mix (1.5 µl R1 (5’-CTACACGACGCTCTTCCGATCT-3’), 1.5 µl TSO (5’-A AGCAGTGGTATCAACGCAGAG-3’) (3µM each, ordered from Eurofins), and 2 µl NfW with following PCR settings:

Step 1: 98°C − 3 min, step 2: 98°C - 20 sec, step 3: 65°C - 30 sec, step 4 - 72°C: 2 min, repeat steps 2 to 4 four times for a total of 5 cycles, step 5: 72°C - 1 min, step 7: 4°C - hold.

Following PCR, a 1.4X SPRI bead cleanup and size selection was performed using Roche Kapa HyperPure Beads (08963835001, Roche) and eluted in 30 µl NfW. Concentration and size distribution were assessed using Qubit 3.0 fluorometer (DNA HS Assay Kit) and TapeStation (HS D5000).

##### Probe hybridization

Probe-based target enrichment of B and T cell receptor (BCR/TCR) sequences was performed using the KAPA Hypercap Workflow V3.5 (Roche), with KAPA HyperCap Target Enrichment Probes, KAPA HyperCapture Bead Kit and KAPA HyperCapture Reagent Kit (Roche). The probe sequences for specific enrichment were kindly provided by Camilla Engblom (Engblom2023 and personal communication).

To hybridize the sample to KAPA HyperCap Target Enrichment Probes, up to 1000 ng of the amplified cDNA in 45 µl NfW was used. To block repetitive regions, 20 µl human COT DNA (included in Roche HyperCapture Reagent Kit) was added. After binding the sample to 130 µl KAPA HyperPure Beads, the bead-bound DNA was eluted in a blocking-oligos-mix for the 10x sequencing adapters (2.5 µl R1, 2.5 µl TSO (100 µM each), and 8.4 µl NfW. Following addition of 43 μl Hybridization Master Mix (according to Roche HyperCapture Reagent Kit) and incubation on a magnet, 56.4 μl eluate was mixed with 4 μl of the KAPA target enrichment probes, prepared according to the manufacturer’s instructions. Hybridization was performed in a thermo cycler with following setting:

Step 1: 95°C – 5 min, step 2: 55°C – 16 to 22 h; Lid: 105°C.

##### Wash and recover of the captured cDNA

After preparation of KAPA HyperCapture Beads, according to the manufacturer’s instructions, the hybridized sample was incubated with the KAPA HyperCapture Beads for 15 min. Beads were washed and bead-bound cDNA was eluted according to the manufacturer’s instructions. Finally, enriched cDNA was eluted in 20 μl NfW.

##### Post-capture amplification

The entire enriched cDNA sample was amplified to increase the final sequencing library, using 25 µl of KAPA HiFi HotStart Ready Mix (2X) (Roche), and 5 µl of 10x cDNA primer mix (2.5 µl R1, 2.5 µl TSO (20 µM each), with following PCR settings:

Step 1: 98°C - 3 min, step 2: 98°C - 20 sec, step 3: 65°C - 30 sec, step 4 - 72°C: 2 min, repeat steps 2 to 4 nineteen times for a total of 20 cycles, step 6: 72°C - 1 min, step 7: 4°C - hold.

Following PCR, a 1.4X SPRI bead cleanup and size selection was performed using Roche Kapa HyperPure Beads (08963835001, Roche) and library was eluted in 20 µl NfW. Concentration and size distribution were assessed using Qubit 3.0 fluorometer (DNA HS Assay Kit) and TapeStation (HS D5000).

##### ONT Sequencing and base-calling

AIRR libraries were prepared for ONT sequencing using the Ligation Sequencing Kit V14 (SQK-LSK114, ONT) according to the manufacturer’s protocol, using 300 fmol input cDNA. Concentration and size distribution were assessed using Qubit 3.0 fluorometer (DNA HS Assay Kit) and TapeStation (HS D5000). Individual samples with a final loading amount of 35 fmol were sequenced on single-MinION flow cells R10.4.1 according to the manufacturer’s protocol. Sequencing output was saved in pod5 format. ONT basecalling was performed using *dorado duplex* (v0.9.1) integrated in *longairr basecall* using the super accurate (sup) model (dna_r10.4.1_e8.2_400bps_sup@v5.0.0) and a minimal quality filter threshold of 5 (*--minqscore 5*). Basecalling was performed using a NVIDIA GeForce RTX 3060.

#### PacBio sequencing of spatial AIRR library

PacBio sequencing of the final AIRR library sample was performed by Novogene’s sequencing service (Novogene, China). Briefly, after cDNA quality control, the SMRTbell library was generated by ligating hairpin-shaped sequencing adapters to both ends of the DNA fragments. Incompletely ligated or failed products were removed using exonuclease treatment. Sequencing primer were annealed to the SMRTbell templates, followed by binding of the sequence polymerase to the annealed templates. After final library quality control, the library was sequenced on one PacBio Revio SMRT Cell (v13.1). Reads shorter than 50 nt were filtered out by the company during read processing. Adapter filtering followed default parameters as stated by the company.

#### ONT sequencing of bulk AIRR library

For sequencing of immunoglobulin heavy chain transcript repertoires from bulk tumor tissue, cryopreserved glioma tissue blocks embedded in OCT Tissue Tek were used. Total RNA was extracted from 10 x 10 μm fresh frozen tissue sections using the RNeasy Mini Kit according to the manufacturer’s protocol “Purification of Total RNA from Animal Tissues” (Qiagen, Germany). RNA concentration was determined using a Qubit 3.0 Fluorometer using the Qubit RNA HS Assay Kit (Thermo Fischer Scientific, United States). The RNA concentration and quality were assessed using Qubit 3.0 fluorometer (RNA HS Assay Kit) and TapeStation (HS RNA).

cDNA synthesis and immunoglobulin heavy chain cDNA amplification were performed using the SMART-Seq Human BCR (with UMIs) kit (Takara Bio Inc, United States) according to the manufacturer protocol. cDNA concentration and size distribution were assessed using Qubit 3.0 (DNA HS Assay Kit) and TapeStation (HS D5000).

Amplicons were prepared for long-read ONT sequencing using the Ligation Sequencing Kit V14 (SQK-LSK114, ONT) according to the manufacturer’s protocol, using 200 fmol input. Concentration and size distribution were assessed using Qubit 3.0 fluorometer (DNA HS Assay Kit) and TapeStation (HS D5000). Individual samples with a final loading amount of 20 fmol were sequenced on single-MinION flow cells R10.4.1 according to the manufacturer protocol (ONT). Basecalling was performed using dorado (ONT) as described for spatial AIRR ONT basecalling.

### COMPUTATIONAL METHODS

#### LongAIRR software implementation

*LongAIRR* is organized into six modular components that together implement a complete workflow for long-read AIRR data processing (**Figure 1b**). The design emphasizes modularity, interoperability with established AIRR analysis frameworks, and scalability for large long-read datasets. The modules cover **(1)** basecalling of ONT raw reads, **(2)** length- and quality filtering, **(3)** demultiplexing of multiplexed bulk sequencing runs, **(4)** UMI and SPBC annotation including sequence group based alignment and adaptive filtering followed by consensus building, **(5)** optional sequence segment annotation such as constant region assignment, and **(6)** V(D)J allele annotation using reference databases (Lefranc2005, Ye2013).

The software modules are implemented in bash and integrate dedicated Python scripts as well as established third-party tools, including components of the Immcantation framework (Gupta2015, VanderHeiden2014). Modules can be executed independently or combined into a streamlined pipeline using workflow managers such as Snakemake (Mölder2021).

All modules generate standardized intermediate outputs (FASTA / FASTQ) and AIRR-compliant TSV files. *LongAIRR* extends this output schema with additional metadata columns capturing UMI and SPBC annotations, as well as molecule-level grouping information such as original and retained read counts following filtering and consensus building. This extended schema preserves full compatibility with AIRR Community standards (VanderHeiden2018) while retaining information necessary for spatial- and UMI-aware downstream AIRR analyses.

##### LongAIRR Pre-processing Modules

###### (1) ONT Basecalling (only for ONT raw data)

The first module, *longairr basecall*, integrates the ONT *Dorado duplex basecaller* (ONTDoradoGit) to process raw sequencing reads in ONT’s pod5 format (**Figure 1b**, **Module 1**). *Dorado* generates duplex reads, in which both complementary DNA strands of a molecule are sequenced, as well as simplex reads derived form only a single strand. Read classification is encoded in the *dx* BAM tag: *dx:1* denotes a duplex read, *dx:0* denotes a simplex read without duplex offspring, and *dx:-1* denotes a simplex read for which a corresponding duplex read was successfully generated and is part of *dx:1* (ONTDoradoGit, ONTDoradoDoc). Duplex rates vary across libraries and sequencing conditions and are typically low, often below 10% (AjaMacaya2025, Sanderson2023). Owing to the combined strand information, duplex reads exhibit improved basecalling accuracy compared to simplex reads (Hall2024, ONTDoradoGit, ONTDoradoDoc).

*LongAIRR* allows users to retain duplex reads only (*dx:1*), simplex reads only (*dx:0*), or a combined dataset (*dx:1* and *dx:0*). To avoid redundancy, reads labeled *dx:-1* are excluded from downstream processing. Selected reads are converted into FASTQ or FASTA format using *samtools view* and *samtools bam2fq* for subsequent analysis (Danecek2021, Li2009). Quality summaries for the basecalled raw reads are generated with *NanoPlot* (DeCoster2023), providing an overview of read length and quality distributions prior to filtering.

###### (2) Sequence length and quality filtering

The second module, *longairr filter*, takes FASTQ files (from module 1 for ONT data or from basecalled PacBio data) as an input and applies customizable quality and length thresholds in order to retain high-quality reads and reduce computational load in subsequent steps (**Figure 1b**, **Module 2**). In contrast to the basecalling module, this step works platform-agnostic and can process FASTQ files generated by ONT as well as HiFi reads from PacBio sequencing.

Filtering is performed using *seqkit seq* and *seqkit fq2fa* (Shen2016, Shen2024). Quality summaries of the filtered data are again generated using *NanoPlot* (DeCoster2023), providing an overview of read length and quality distributions after filtering.

###### (3) Sample demultiplexing

For multiplexed bulk libraries, *LongAIRR* can perform demultiplexing based on unique dual index (UDI) sequences provided as an additional FASTQ input file (**Figure 1b**, **Module 3**). Sample demultiplexing is implemented using the *Immcantation presto* scripts *MaskPrimers* and *SplitSeq*, generating sample-specific FASTA output suitable for downstream analysis (VanderHeiden2014).

To ensure robust index assignment, orientation-aware processing is applied, and UDI sequences are mapped across both read orientations. The resulting sample-resolved outputs enable parallel processing of multiple samples derived from a single sequencing run.

##### LongAIRR Annotation Modules

###### (4) Annotation of UMIs and SPBCs, Adaptive Filtering and Consensus Building

UMIs are short nucelotide barcodes incorporated during library preparation that enable molecule-level error correction in consensus sequences (Kivioja2012). In spatial transcriptomics datasets generated using Visium V1 or Visium HD 3′, sequencing reads additionally contain spot-specific or grid-based SPBCs that encode spatial origin in the tissue section.

The fourth module, l*ongairr collapse*, performs annotation of UMIs and, for spatial datasets, SPBCs from Visium V1 or Visium HD 3′ libraries (**Figure 1b**, **Module 4**). Annotation requires a FASTA file containing a user-defined UMI anchor sequence, and for spatial datasets, a reference list of valid spatial barcodes provided by 10x Genomics (10xSpaceRanger). Reads containing invalid SPBCs are discarded, or in case mismatches are permitted, the closest valid SPBC from the provided reference list is assigned. Anchor matching is performed using *Immcantation presto’s MaskPrimers* (VanderHeiden2014). For datasets that were not processed with *longairr demux* (**Figure 1b**, **Module 3**), both read orientations are evaluated during anchor detection to ensure robust barcode recovery independent of strand direction.

Following UMI and SPBC annotation, reads are grouped by UMI (bulk) or combined UMI-SPBC identifier (spatial), with each group representing an individual captured molecule. This ensures that molecules originating from different spatial locations within the tissue are processed independently.

Prior to consensus building, two complementary filtering strategies are applied to improve consensus building results. The first approach applies fixed-length thresholds via *seqkit seq* (Shen 2016, Shen204), retaining reads within user-defined bounds expected to capture full-length receptor transcripts. For the second filter approach, *LongAIRR* implements an adaptive filtering strategy applied to individual UMI-SPBC groups above a specified minimum size (parameter *filter-min-size*, default value 5). In this step, read-length distributions are evaluated per group, and only reads falling within a defined margin around the group-specific dominant peak length (i.e., the most populated sequence length bin) are retained. This approach preserves the predominant full-length transcript representation by filtering relative to the overall length distribution within each UMI-SPBC each group.

To enable scalable processing of large UMI-SPBC groups, *longairr collapse* additionally applies a subsampling strategy prior to multiple sequence alignment (MSA). Groups exceeding a user-defined size threshold (parameter *n-subsample*, default value 750) are reduced to manageable subsets before alignment using *presto’s AlignSets* (MUSCLE 3.8) (Edgar2004, VanderHeiden2014). Following group-specific MSA, reads are partitioned into chunks that preserve complete groups, enabling parallel consensus generation with *BuildConsensus* (VanderHeiden2014). Users may configure subsample size, number of groups per chunk, and the level of parallelization to match available computational resources. Read-specific metadata including the annotated UMI and SPBC sequences, as well as original-, filtered- and subsampled group-sizes are annotated in each read’s FASTA header, supporting traceability throughout downstream analysis (**Figure 1b**, **Module 4**).

###### (5) Additional read-segment annotation

The *longairr seqtag* module provides flexible anchor-based annotation of additional read segments. For instance, in immunoglobulin-based bulk datasets, constant region anchor can be used to assign isotype-families. In combined immunoglobulin and T cell receptor spatial AIRR libraries, the module enables separation of sequences into locus-specific subsets prior to V(D)J annotation (**Figure 1b**, **Module 5**).

Anchor sequences are user-defined input provided in FASTA format and are mapped using *presto’s MaskPrimers* and *SplitSeq* (VanderHeiden2014), which allows separation of sequences into multiple files based on user-defined regular expression, e.g., identifiers associated with provided anchor-sequences.

###### (6) V(D)J gene annotation

The final module, *longairr airr*, performs loci-specific annotation of V(D)J and constant gene segments using *Change-O’s AssignGenes igblast* script with user-specified IMGT germline references (Gupta2015, Lefranc2005, Ye2013), which can be downloaded and configured during *LongAIRR’s* installation (**Figure 1b**, **Module 6**). Annotated reads are transformed into tabular, AIRR-compliant outputs using *Change-O’s MakeDb* and *ParseDb* scripts (Gupta2015).

#### *LongAIRR* data processing

All datasets generated in this study were processed using *LongAIRR*. Basecalling of ONT datasets was performed using *longairr basecall*, as described in this study. PacBio datasets consisted of HiFi reads with minimal length of 50 nt and were used directly for *LongAIRR* processing as received by the company. Following ONT basecalling, all datasets were processed using a custom Snakemake workflow consisting of the *LongAIRR* modules: *longairr filter*, *longairr demux* (bulk datasets only), *longairr collapse*, *longairr seqtag*, and *longairr airr* on an HPC cluster. During the *longairr filter* step, reads were filtered using a minimum read length of 420 nt, maximum read length of 6,000 nt, and a minimum PHRED quality threshold of Q10 (S1_ONT_Q10) or Q17 for all other datasets (ONT Visium V1, PacBio Visium V1, ONT Visium HD 3′, ONT Bulk, see **Table 1**).

Filtered reads were next processed using *longairr collapse*. After UMI and SPBC annotation for spatial datasets, reads were targeted length-filtered with a minimum length of 800 nt and a maximum length of 3,800 nt. For UMI-SPBC groups containing more than four reads, adaptive length filtering was applied by retaining reads within 5% of the most abundant read length within the sequence group prior to multiple sequence alignment and consensus sequence construction. Additionally large UMI-SPBC groups where subsampled to 750 reads per group.

A comprehensive list of all *LongAIRR* parameters used to process the spatial samples can be found in **Table S2**.

#### Sequence quality assessment with PHRED scores

Per-read PHRED quality scores were calculated from FASTQ quality strings by converting base-level quality characters to PHRED scores, transforming these into per-base error probabilities, averaging the error probabilities across the read, and converting the resulting mean error probability back to a PHRED score. Dataset-level mean read quality values, as reported in Fig. 2b-2d, were then calculated from per-read PHRED scores using the same error-probability-based averaging scheme implemented in NanoPlot (DeCoster2023).

#### Subsampling consensus analysis

To evaluate the robustness of consensus generation across different UMI-SPBC group sizes, representative large groups were selected from ONT and PacBio datasets and reprocessed using the internal *LongAIRR* consensus-building logic across a range of subsample sizes. For each selected UMI-SPBC group, reads were first filtered by length (800-3,800 nt) and then subsampled to predefined group sizes. For each subsample size, five independent replicates were generated using fixed random seeds (1-5). Groups containing at least five reads were additionally subjected to adaptive length filtering using the same parameters applied in the *LongAIRR* workflow (peak margin 5% around the dominant read-length bin), followed by multiple sequence alignment and consensus construction.

Consensus sequences generated from each subsample were compared to the corresponding consensus derived from the full group which was used as the ground-truth reference. Sequence similarity was assessed for the variable receptor region spanning FWR1-FWR4, while flanking adaptor and constant regions were excluded. Similarity was quantified as Hamming distance and percent identity after pairwise alignment. Productive status was additionally assessed based on *IgBLAST* (Ye2013) of the resulting consensus sequences.

#### Rarefaction analysis

Rarefaction analysis was performed to assess the relationship between sequencing depth and the number of recovered productive spatial AIRR UMI-SPBC groups. Raw FASTQ files from the library sequenced with ONT and PacBio (S1_ONT, S1_PB) were subsampled at predefined fractions using *seqkit sample* (Shen2016, Shen2024), generating subsets containing 0.1%, 1%, 2.1%, 4.6%, 10%, 21.5%, 46.4%, 60%, 75%, 90% and 100% of the original reads. Subsampling was performed with a fixed random seed. Each subsampled dataset was processed independently using the *LongAIRR* workflow with the same parameters applied to the full dataset. For the ONT dataset, two different quality filtering strategies were applied which resulted in two rarefaction curves (S1_ONT_Q10 and S1_ONT_Q17).

For each dataset and subsampling level, the number of recovered molecules was determined by counting unique UMI-SPBC groups in the output AIRR tables for both BCR and TCR loci. Groups were further divided according to the number of reads supporting the corresponding consensus sequence: single-read groups (consensus count = 1) and multi-read groups (consensus count > 1) illustrated by the line type in Fig. 3b.

#### Statistical analysis using Cliff’s delta

Differences between V-region identity distributions were quantified using Cliff’s delta (δ), a non-parametric effect size measure that assesses the degree of separation between two distributions. Cliff’s delta was calculated using the *effsize::cliff.delta()* function in R (Torchiano2016). Pairwise comparisons were performed in a directional manner, with the first dataset in each comparison defined as group 1 and the second as group 2. The following comparisons were evaluated: (1) ONT_R9_original vs ONT_R9_longAIRR, (2) ONT_R9_longAIRR vs S3_ONT_Q17, (3) ONT_R9 longAIRR vs S4_ONT_Q17_HD, (4) ONT_R9_original vs S3_ONT_Q17, and (5) ONT_R9_original vs S4_ONT_Q17_HD. Negative Cliff’s delta values indicate higher V-region identity in the second dataset, whereas positive values indicate higher V-region identity in the first dataset of each comparison.

#### Mutual information analysis

Pairwise mutual information between the Ig heavy and light chain clones were computed using the *mutinformation()* function of the *infotheo* package (Meyer2014). Ig heavy chain clones were assigned using the *hierachicalClones()* function of the *scoper* package (Gupta2017). Ig light chain clones were defined by identical V call, J call and junction length.

#### 10x Visium GEX quantification and downstream analyses

Spatial gene expression datasets were generated using the 10x Visium V1 platform described in this study. Gene expression was quantified from the Illumina sequencing output using Space Ranger (v4.0.1) (10xSpaceRanger) and the 10x Genomics refdata-gex-GRCh38-2020-A reference package, which is based on the GRCh38 human genome.

##### Pre-processing and quality control

Each gene in the SpatialExperiment object was annotated with its biotype and chromosomal location using the 10x Genomics GRCh38 reference GTF file (refdata-gex-GRCh38-2020-A). To focus the analysis on relevant transcripts, long non-coding RNA (lncRNA) genes were excluded. The retained biotype set comprised: protein_coding, IG_C_gene, IG_C_pseudogene, IG_D_gene, IG_J_gene, IG_J_pseudogene, IG_V_gene, IG_V_pseudogene, TR_C_gene, TR_D_gene, TR_J_gene, TR_J_pseudogene, TR_V_gene, and TR_V_pseudogene.

Per-spot quality control (QC) metrics were computed using *scuttle::addPerCellQC()* (McCarthy2016) with mitochondrial genes specified as a subset, yielding total UMI counts, number of detected genes, and the percentage of mitochondrial reads per spot. Per-gene QC metrics were computed using *scuttle::addPerFeatureQC()*. Filtering steps included gene-level filtering with genes detected (count > 0) in fewer than 3 spots were removed. Spot-level filtering applied the following thresholds: a minimum of 100 detected genes and a minimum of 100 total UMIs. Mitochondrial filtering was applied conditionally: the 95th percentile of the per-spot mitochondrial percentage was computed as the threshold, but this filter was activated only if at least one spot exceeded 20% mitochondrial content. A spot was removed only if it simultaneously fell below the gene detection threshold, below the UMI threshold, and above the mitochondrial threshold.

Library size factors were computed using *scuttle::computeLibraryFactors()*, and log-normalized counts were generated with *scuttle::logNormCounts()* (McCarthy2016).

##### Cell-Type Signature Scoring with UCell

Cell-type signature scores were computed using the UCell algorithm (v2.12.0) (Andreatta2021). Scoring was performed on the log-normalized expression matrix. The marker gene list for B cells was derived from GBmap-predicted cell type (annotation level 4) (RuizMoreno2025) and is available in Table S1.

##### Tertiary Lymphoid Structure Identification

TLS candidate regions were defined using a B cell aggregate–focused strategy. The B cell UCell score served as the primary spatial signal. Spots were classified as B cell-enriched if their B cell score exceeded the 90th percentile of the score distribution.

Spatial clustering of B cell-enriched spots was performed using the density-based spatial clustering algorithm implemented in the *dbscan* R package (v1.2.2) (Hahsler2019). The algorithm was applied to the spatial coordinates of B cell-enriched spots only. Clustering parameters were set to *eps* = 200, defining the neighbourhood radius in the spatial coordinate system (approximately two Visium spot spacings) and *minPts* = 3. A cluster was confirmed as a TLS region if it satisfied the following criteria simultaneously: *n_spots* ≥ 3, *n_spots* ≤ 1000 and if at least 50% of the spots in the cluster had B cell scores above the 90th percentile. All remaining spots were labeled as non-TLS spots.

#### Figure generation

All plots were generated in R (v4.5.0) using *ggplot2* (Wickham2016). Multi-panel figures were assembled using *patchwork* (Pedersen2019) and *cowplot* (Wilke2025) packages. Fig. 1a, 1b, 4a, 5a where created using Google Slides.

## CODE AND DATA AVAILABILITY

The LongAIRR software is available as an open-source repository on GitHub: https://github.com/AGImkeller/LongAIRR under the MIT License including extensive documentation and an exemplary Snakemake workflow using *LongAIRR,* used for data processing in this study.

The AIRR datasets generated and analyzed in this study, including all config files that inherit all parameters passed to *LongAIRR* in the used snakemake workflow, will be made publicly available through a suitable repository upon publication. GEO Accession numbers for the analyzed public data can be found under **GSM7658223** for the analyzed data and **SRX21134879** for raw data from Benotmane et al (Benotmane2023).

## AUTHOR CONTRIBUTIONS

J.S., S.O.I. and K.I. conceived the project. J.S. developed the *LongAIRR* software. S.O.I., Z.M. and K.I. established the experimental workflow and conducted experiments. K.W. provided sample material and supported the histomorphological and neuropathological annotation. K.I. supervised experiments and software development. J.S., C.G.A., L.M.H. and K.I. performed data analysis. J.S. and K.I. generated figures and wrote the manuscript. All authors reviewed and approved the final manuscript.

## ACKNOWLEDGEMENTS

The authors would like to thank Aakanksha Singh for helpful feedback on the manuscript. The authors thank the UCT tumor documentation (Sandra V. Klein, Andrea Wolf) and the UCT Biobank (Kristina Goetze) for providing samples and advice.

## SUPPLEMENT

### SUPPLEMENTARY FIGURES

**Figure S1:**
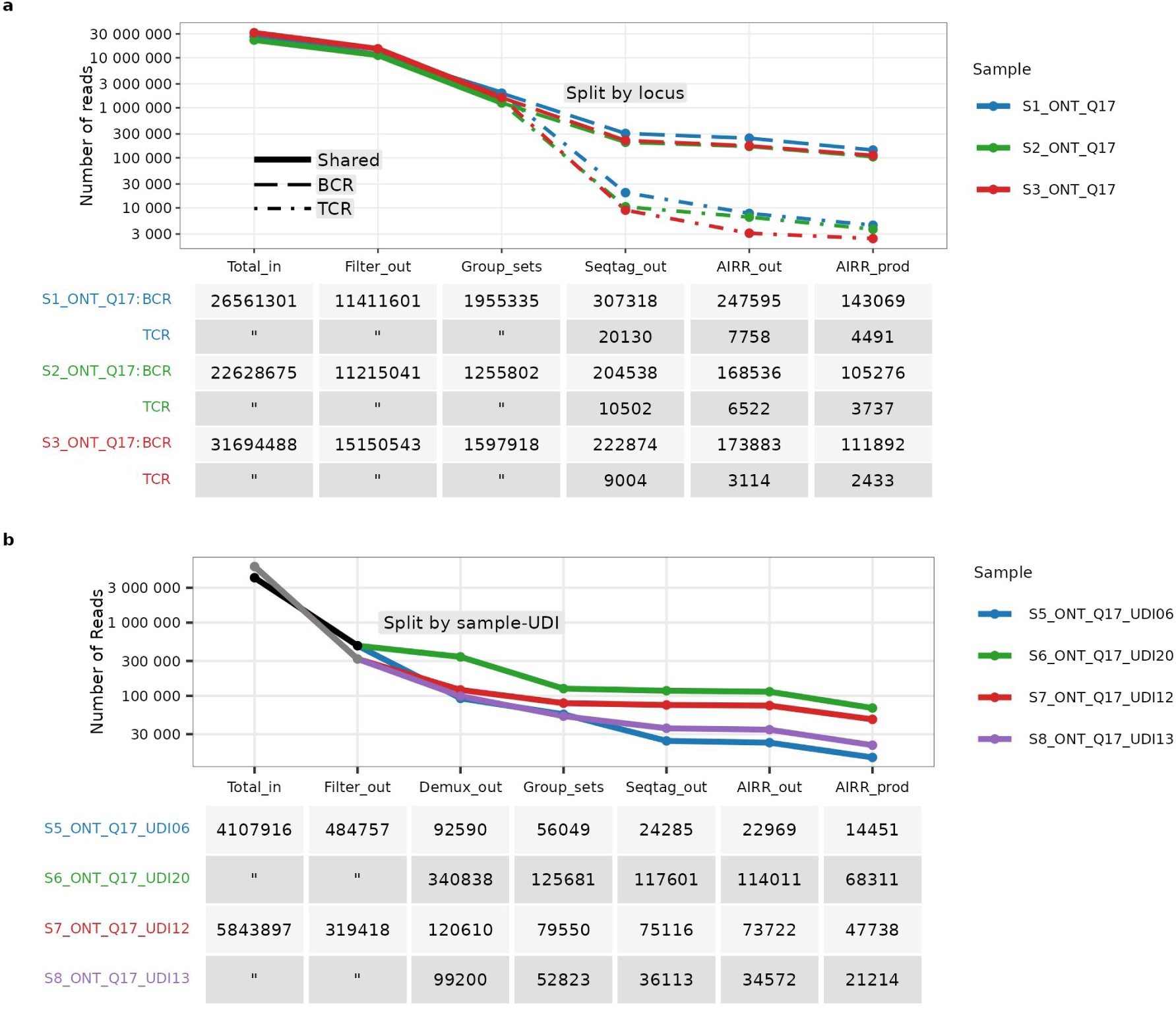
Read retention across *LongAIRR* processing steps for spatial and bulk datasets. **a:** Number of sequences retained across *LongAIRR* processing steps: length- and quality-filtered reads (*Filter_out*), UMI-SPBC group consensus sequences (*Group_sets*), sequences with detected BCR or TCR constant sequence segments (*Seqtag_out*) and V(D)J-annotated AIRR sequences (all: *AIRR_out*; productive: *AIRR_prod*). Colors indicate datasets (see Table 1). Line types indicate loci: combined BCR and TCR reads (solid), BCR (large dashes) and TCR (small dashes) after constant region annotation and BCR/TCR splitting. **b:** Similar analysis and representation as in (a) illustrating read retention for bulk BCR sequencing datasets. Grey and black line segments indicate combined reads prior to demultiplexing, after which reads are separated into sample-specific datasets. Samples S5_ONT_Q17_UDI06 and S6_ONT_Q17_UDI20 (black) were multiplexed and sequenced together, as were S7_ONT_Q17_UDI12 and S8_ONT_Q17_UDI13 (grey).

**Figure S2:**
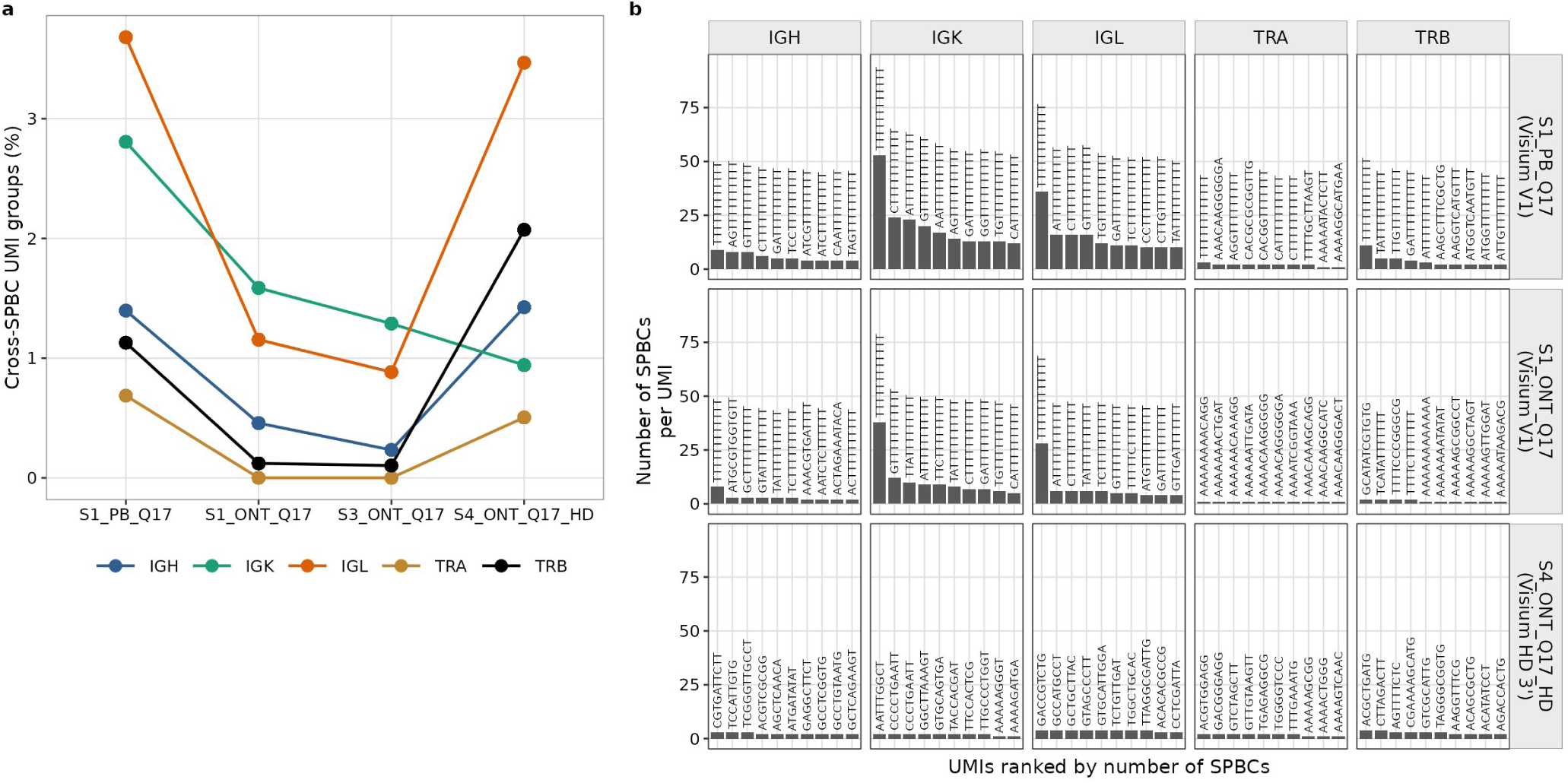
Multi-SPBC UMI groups across sequencing platforms. **a:** Percentage of UMI groups associated with multiple SPBCs across datasets (S1_PB_Q17, S1_ONT_Q17, S3_ONT_Q17, S4_ONT_Q17_HD). Colors indicate transcript type (Ig heavy, Ig κ, Ig λ, TCR α, TCR β). **b:** Top 10 UMI groups with the highest number of associated SPBCs, shown per sequencing platform andtranscript type (Ig heavy, Ig κ, Ig λ, TCR α, TCR β). Rows correspond to datasets (S1_PB_Q17, S1_ONT_Q17, S4_ONT_Q17_HD). Bars represent individual UMI groups, with UMI sequences indicated as labels.

**Figure S3:**
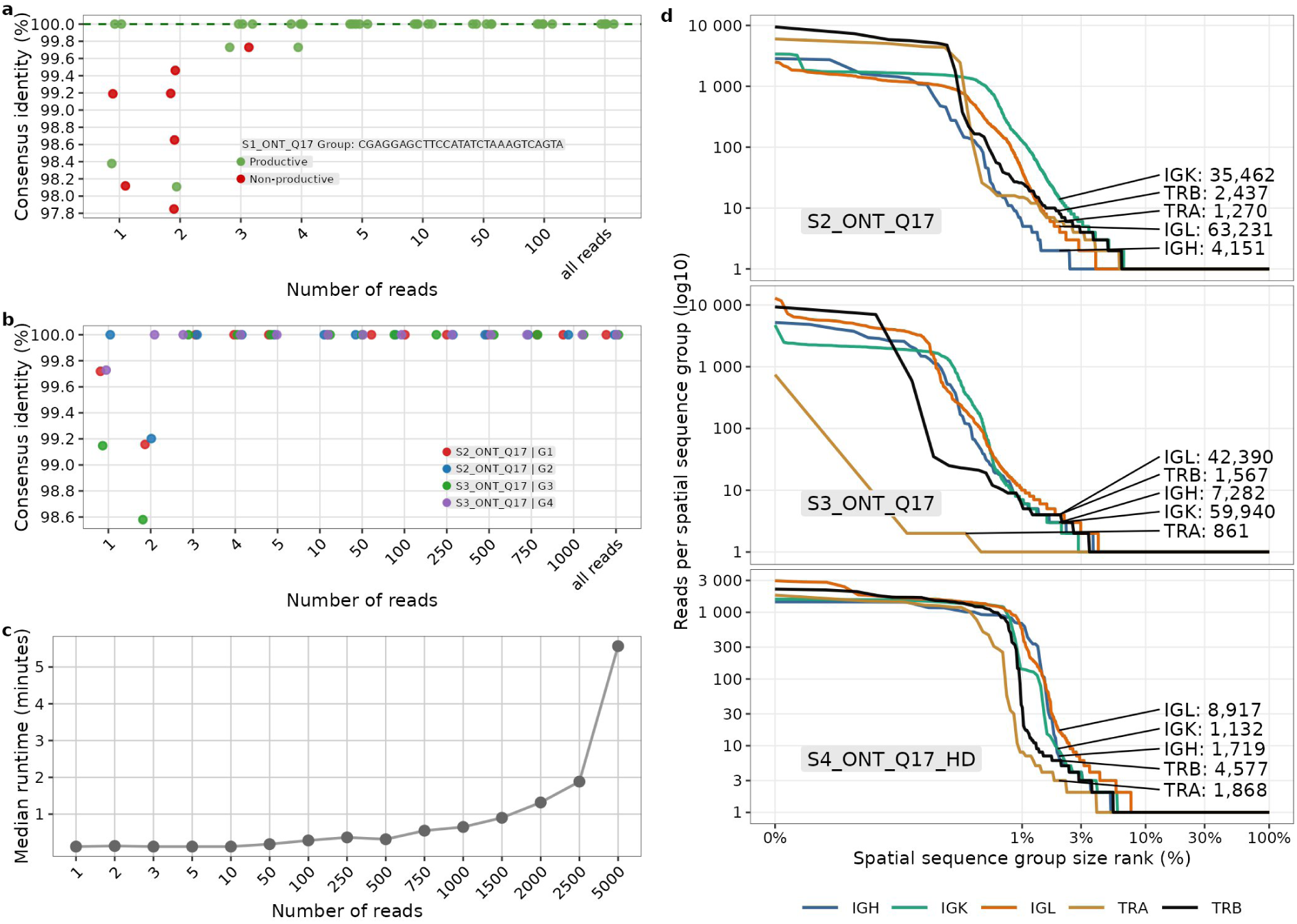
Subsampling-based assessment of consensus sequence robustness, runtime and UMI group size distributions. **a:** Subsampling analysis of consensus sequence robustness for a representative group from S1_ONT_Q17, corresponding to the group shown in Fig. 4e (S1_PB_Q17). Consensus sequences from subsampled groups of increasing size are compared to the corresponding full-group consensus. Sequence identity is assessed as hamming distance within FWR1-FWR4 region. UMI-SPBC group identifiers is indicated in the panel. Colors indicate productivity status of the consensus receptor sequence: productive (green); non-productive (red). **b:** Subsampling analysis of consensus sequence robustness for additional representative groups from S2_ONT_Q17 (G1, G2) and S3_ONT_Q17 (G3, G4). Consensus sequences from subsampled groups of increasing size are compared to the corresponding full-group consensus. Points show median identity across replicates (n = 5) and are colored by group. **c:** Runtime of consensus sequence generation across subsample sizes for the representative group shown in Fig. 4d. Points show median runtime across replicates (n = 5). **d:** Number of reads per UMI-SPBC consensus group for additional datasets (S2_ONT_Q17, S3_ONT_Q17, S4_ONT_Q17_HD). Consensus groups are ranked by size on a percentile scale (non-linear spacing on x-axis). Numbers indicate the total number of consensus groups per transcript type (Ig heavy, Ig κ, Ig λ, TCR α, TCR β).

**Figure S4:**
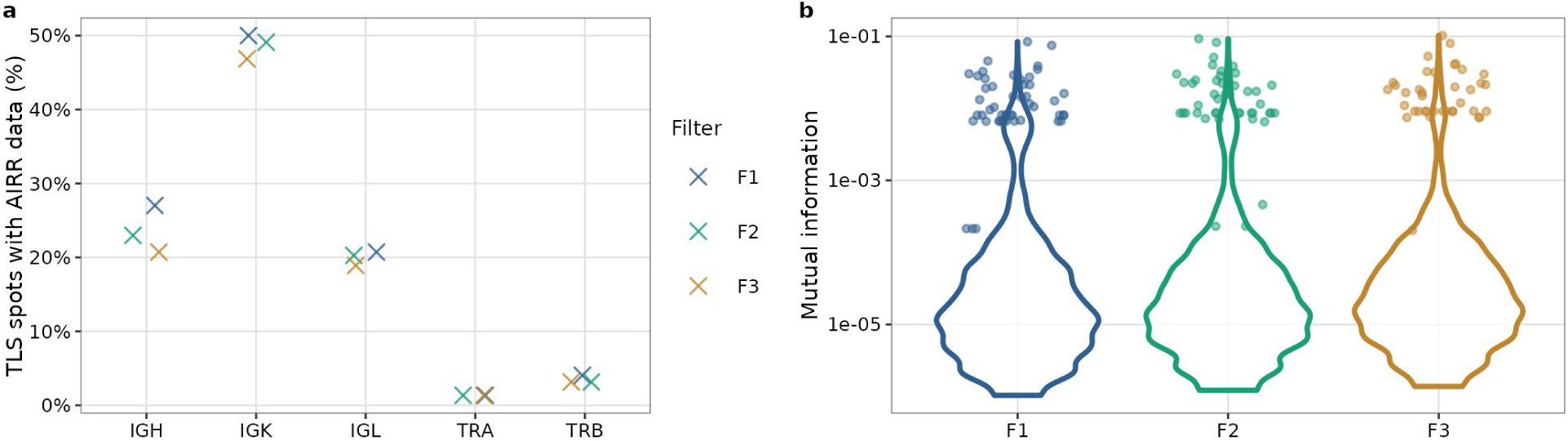
TLS-associated AIRR signal and chain pairing information across downstream filtering strategies. **a:** Relative proportion of TLS spots containing annotated AIRR sequences across transcript types (Ig heavy, Ig κ, Ig λ, TCR α, TCR β) under different filtering strategies (F1-F3). TLS spots are defined based on gene expression profiles (see Methods). **b:** Distribution of mutual information for heavy-light chain pairings across filtering strategies (F1-F3). Points indicate for each heavy chain clone, the pairing with the highest mutual information.

### SUPPLEMENTARY TABLES

**Table S1:**
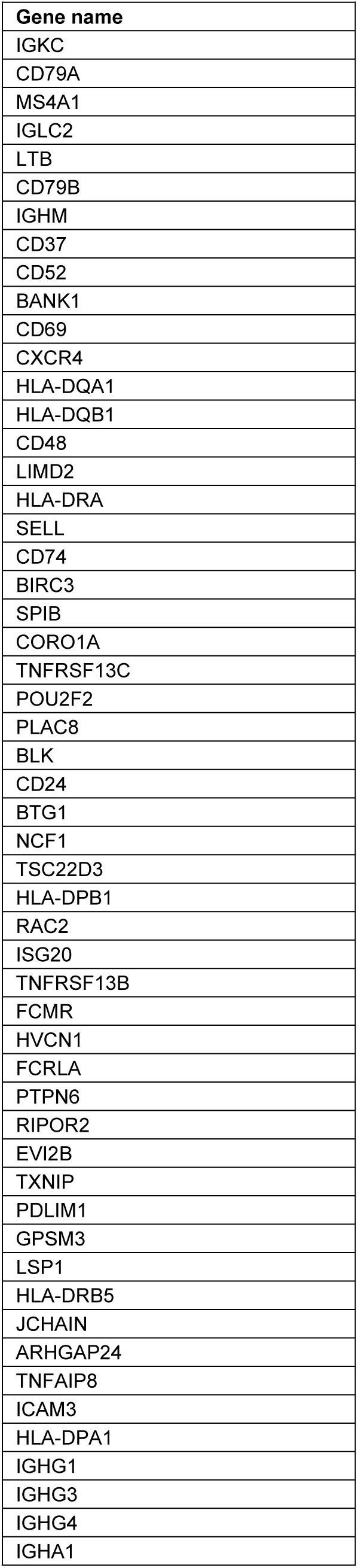
B cell marker gene list used for TLS annotation.

**Table S2:**
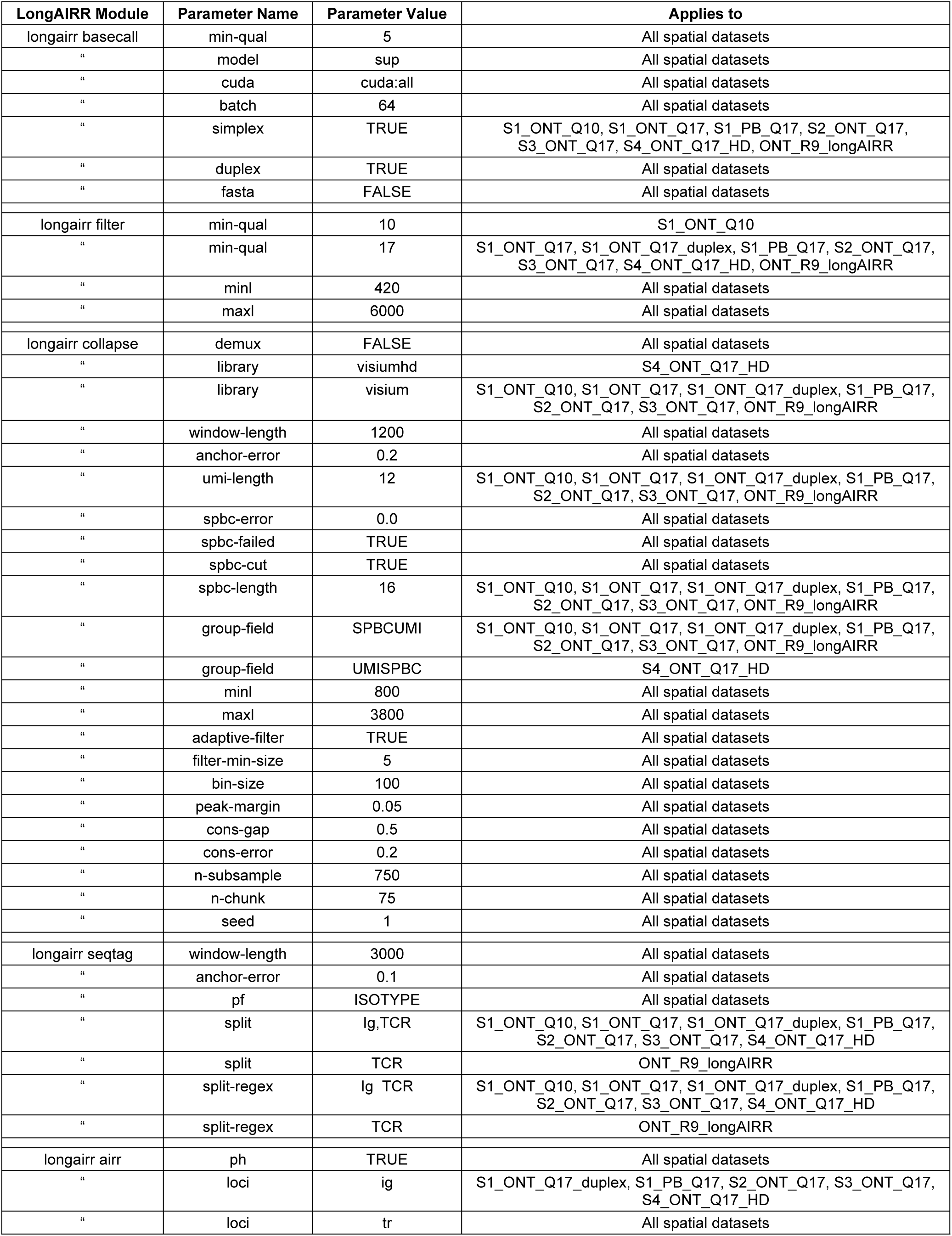
LongAIRR parameters used for processing spatial AIRR datasets. Spatial AIRR datasets correspond to the subset of datasets in Table 1 generated using Visium V1 and Visium HD 3’-based spatial transcriptomics (S1-S4 and ONT_R9_longAIRR). Bulk datasets (S5-S8) were processed separately and are not included in this table.

