## supplement for "Consistent consensus-based annotation of spatial adaptive immune receptor repertoires from long-read sequencing using *LongAIRR*"

### SUPPLEMENTARY FIGURES

a

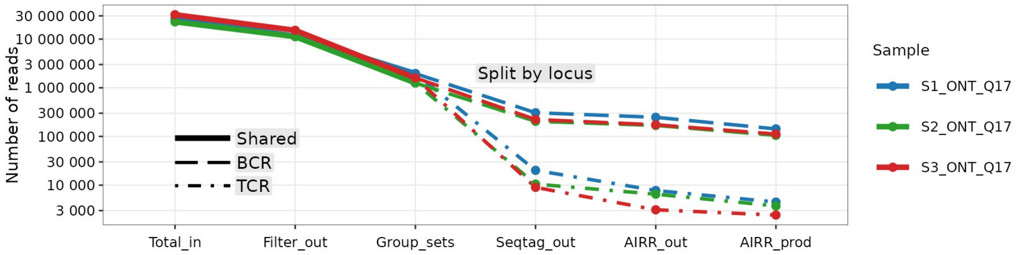

| | Total_in | Filter_out | Group_sets | Seqtag_out | AIRR_out | AIRR_prod |
|---|---|---|---|---|---|---|
| S1_ONT_Q17: BCR | 26561301 | 11411601 | 1955335 | 307318 | 247595 | 143069 |
| TCR | " | " | " | 20130 | 7758 | 4491 |
| S2_ONT_Q17: BCR | 22628675 | 11215041 | 1255802 | 204538 | 168536 | 105276 |
| TCR | " | " | " | 10502 | 6522 | 3737 |
| S3_ONT_Q17: BCR | 31694488 | 15150543 | 1597918 | 222874 | 173883 | 111892 |
| TCR | " | " | " | 9004 | 3114 | 2433 |

b

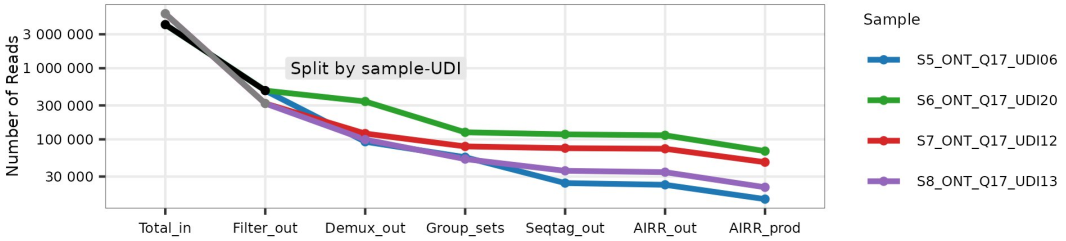

| | Total_in | Filter_out | Demux_out | Group_sets | Seqtag_out | AIRR_out | AIRR_prod |
|---|---|---|---|---|---|---|---|
| S5_ONT_Q17_UDI06 | 4107916 | 484757 | 92590 | 56049 | 24285 | 22969 | 14451 |
| S6_ONT_Q17_UDI20 | " | " | 340838 | 125681 | 117601 | 114011 | 68311 |
| S7_ONT_Q17_UDI12 | 5843897 | 319418 | 120610 | 79550 | 75116 | 73722 | 47738 |
| S8_ONT_Q17_UDI13 | " | " | 99200 | 52823 | 36113 | 34572 | 21214 |

**Figure S1: Read retention across *LongAIRR* processing steps for spatial and bulk datasets**

**a:** Number of sequences retained across *LongAIRR* processing steps: length- and quality-filtered reads (*Filter_out*), UMI-SPBC group consensus sequences (*Group_sets*), sequences with detected BCR or TCR constant sequence segments (*Seqtag_out*) and V(D)J-annotated AIRR sequences (all: *AIRR_out*; productive: *AIRR_prod*). Colors indicate datasets (see Table 1). Line types indicate loci: combined BCR and TCR reads (solid), BCR (large dashes) and TCR (small dashes) after constant region annotation and BCR/TCR splitting.

**b:** Similar analysis and representation as in (a) illustrating read retention for bulk BCR sequencing datasets. Grey and black line segments indicate combined reads prior to demultiplexing, after which reads are separated into sample-specific datasets. Samples S5_ONT_Q17_UDI06 and S6_ONT_Q17_UDI20 (black) were multiplexed and sequenced together, as were S7_ONT_Q17_UDI12 and S8_ONT_Q17_UDI13 (grey).

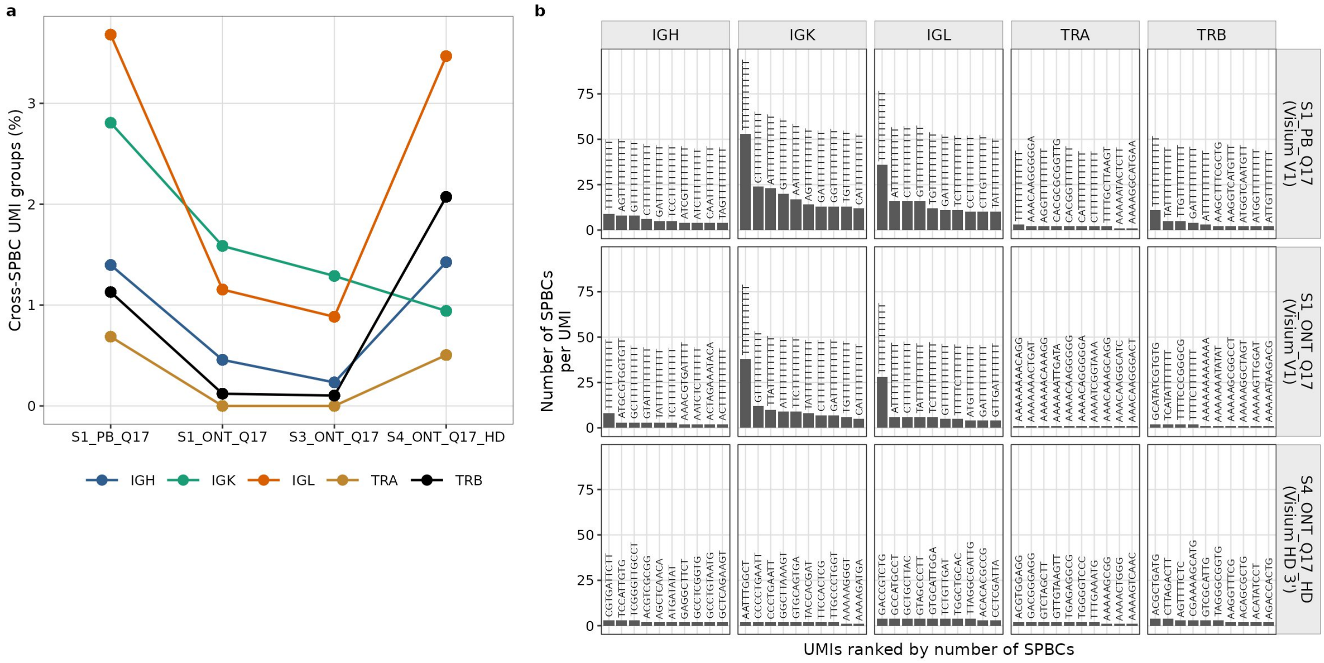

**Figure S2: Multi-SPBC UMI groups across sequencing platforms**

**a:** Percentage of UMI groups associated with multiple SPBCs across datasets (S1_PB_Q17, S1_ONT_Q17, S3_ONT_Q17, S4_ONT_Q17_HD). Colors indicate transcript type (Ig heavy, Ig κ, Ig λ, TCR α, TCR β).

**b:** Top 10 UMI groups with the highest number of associated SPBCs, shown per sequencing platform andtranscript type (Ig heavy, Ig κ, Ig λ, TCR α, TCR β). Rows correspond to datasets (S1_PB_Q17, S1_ONT_Q17, S4_ONT_Q17_HD). Bars represent individual UMI groups, with UMI sequences indicated as labels.

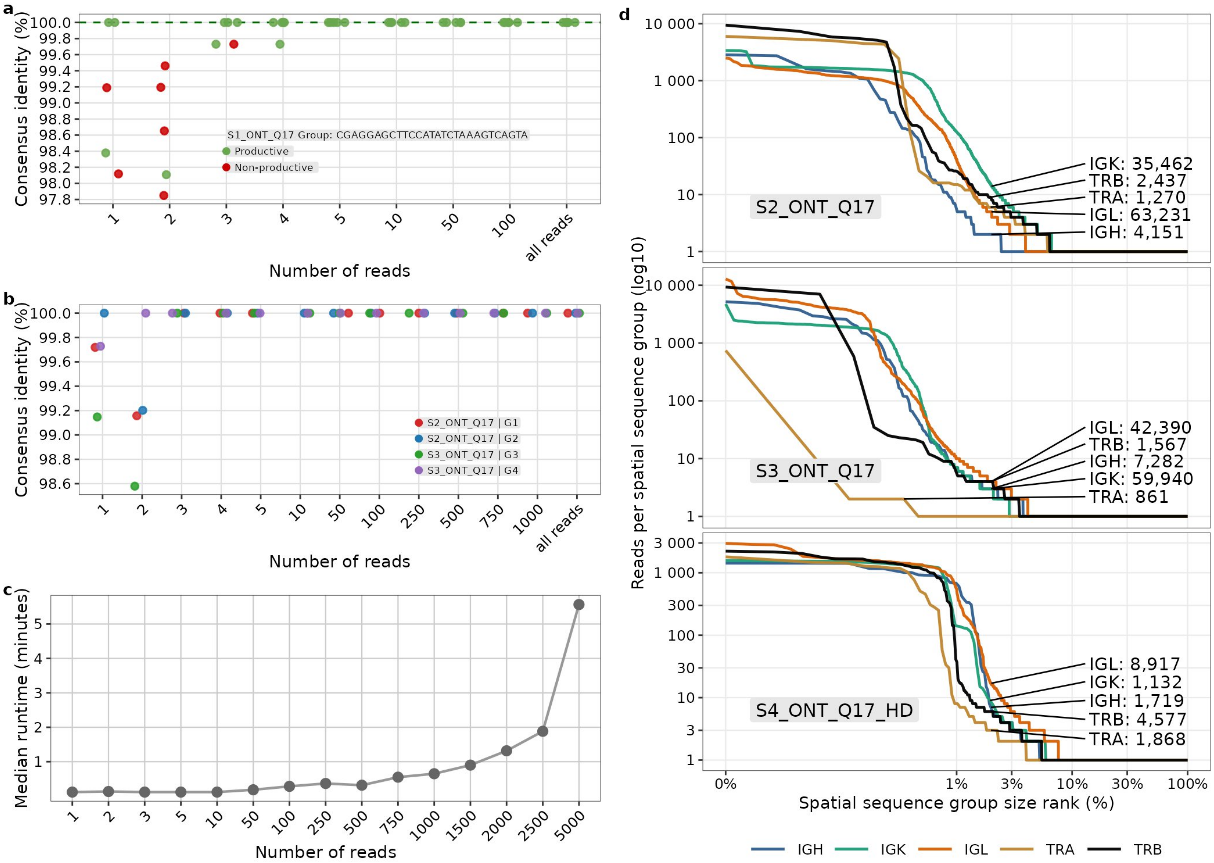

**Figure S3: Subsampling-based assessment of consensus sequence robustness, runtime and UMI group size distributions**

**a:** Subsampling analysis of consensus sequence robustness for a representative group from S1_ONT_Q17, corresponding to the group shown in Fig. 4e (S1_PB_Q17). Consensus sequences from subsampled groups of increasing size are compared to the corresponding full-group consensus. Sequence identity is assessed as hamming distance within FWR1-FWR4 region. UMI-SPBC group identifiers is indicated in the panel. Colors indicate productivity status of the consensus receptor sequence: productive (green); non-productive (red).

**b:** Subsampling analysis of consensus sequence robustness for additional representative groups from S2_ONT_Q17 (G1, G2) and S3_ONT_Q17 (G3, G4). Consensus sequences from subsampled groups of increasing size are compared to the corresponding full-group consensus. Points show median identity across replicates (n = 5) and are colored by group.

**c:** Runtime of consensus sequence generation across subsample sizes for the representative group shown in Fig. 4d. Points show median runtime across replicates (n = 5).

**d:** Number of reads per UMI-SPBC consensus group for additional datasets (S2_ONT_Q17, S3_ONT_Q17, S4_ONT_Q17_HD). Consensus groups are ranked by size on a percentile scale (non-linear spacing on x-axis). Numbers indicate the total number of consensus groups per transcript type (Ig heavy, Ig κ, Ig λ, TCR α, TCR β).

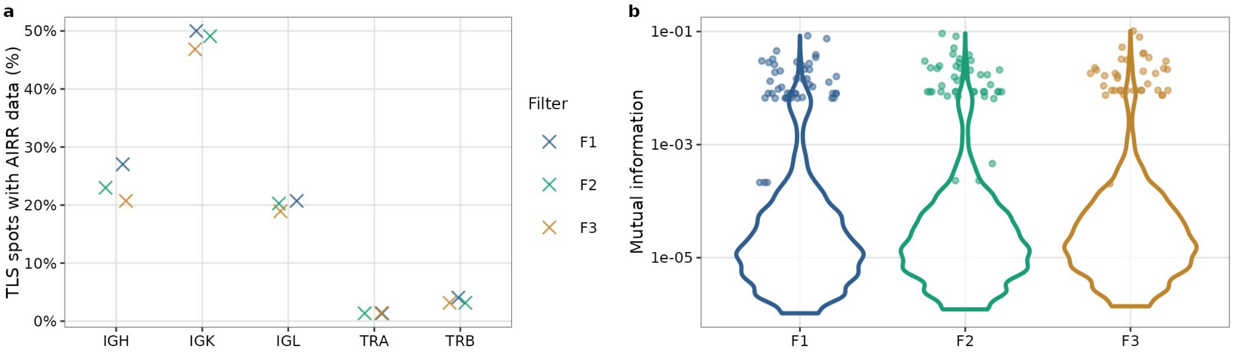

**Figure S4: TLS-associated AIRR signal and chain pairing information across downstream filtering strategies**

**a:** Relative proportion of TLS spots containing annotated AIRR sequences across transcript types (Ig heavy, Ig κ, Ig λ, TCR α, TCR β) under different filtering strategies (F1-F3). TLS spots are defined based on gene expression profiles (see Methods).

**b:** Distribution of mutual information for heavy-light chain pairings across filtering strategies (F1-F3). Points indicate for each heavy chain clone, the pairing with the highest mutual information.

### SUPPLEMENTARY TABLES

**Table S1: B cell marker gene list used for TLS annotation**

| Gene name |
|---|
| IGKC |
| CD79A |
| MS4A1 |
| IGLC2 |
| LTB |
| CD79B |
| IGHM |
| CD37 |
| CD52 |
| BANK1 |
| CD69 |
| CXCR4 |
| HLA-DQA1 |
| HLA-DQB1 |
| CD48 |
| LIMD2 |
| HLA-DRA |
| SELL |
| CD74 |
| BIRC3 |
| SPIB |
| CORO1A |
| TNFRSF13C |
| POU2F2 |
| PLAC8 |
| BLK |
| CD24 |
| BTG1 |
| NCF1 |
| TSC22D3 |
| HLA-DPB1 |
| RAC2 |
| ISG20 |
| TNFRSF13B |
| FCMR |
| HVCN1 |
| FCRLA |
| PTPN6 |
| RIPOR2 |
| EVI2B |
| TXNIP |
| PDLIM1 |
| GPSM3 |
| LSP1 |
| HLA-DRB5 |
| JCHAIN |
| ARHGAP24 |
| TNFAIP8 |
| ICAM3 |
| HLA-DPA1 |
| IGHG1 |
| IGHG3 |
| IGHG4 |
| IGHA1 |

**Table S2: LongAIRR parameters used for processing spatial AIRR datasets**

Spatial AIRR datasets correspond to the subset of datasets in Table 1 generated using Visium V1 and Visium HD 3'-based spatial transcriptomics (S1-S4 and ONT_R9_longAIRR). Bulk datasets (S5-S8) were processed separately and are not included in this table.

| LongAIRR Module | Parameter Name | Parameter Value | Applies to |
|---|---|---|---|
| longairr basecall | min-qual | 5 | All spatial datasets |
| " | model | sup | All spatial datasets |
| " | cuda | cuda:all | All spatial datasets |
| " | batch | 64 | All spatial datasets |
| " | simplex | TRUE | S1_ONT_Q10, S1_ONT_Q17, S1_PB_Q17, S2_ONT_Q17, S3_ONT_Q17, S4_ONT_Q17_HD, ONT_R9_longAIRR |
| " | duplex | TRUE | All spatial datasets |
| " | fasta | FALSE | All spatial datasets |
| longairr filter | min-qual | 10 | S1_ONT_Q10 |
| " | min-qual | 17 | S1_ONT_Q17, S1_ONT_Q17_duplex, S1_PB_Q17, S2_ONT_Q17, S3_ONT_Q17, S4_ONT_Q17_HD, ONT_R9_longAIRR |
| " | minl | 420 | All spatial datasets |
| " | maxl | 6000 | All spatial datasets |
| longairr collapse | demux | FALSE | All spatial datasets |
| " | library | visiumhd | S4_ONT_Q17_HD |
| " | library | visium | S1_ONT_Q10, S1_ONT_Q17, S1_ONT_Q17_duplex, S1_PB_Q17, S2_ONT_Q17, S3_ONT_Q17, ONT_R9_longAIRR |
| " | window-length | 1200 | All spatial datasets |
| " | anchor-error | 0.2 | All spatial datasets |
| " | umi-length | 12 | S1_ONT_Q10, S1_ONT_Q17, S1_ONT_Q17_duplex, S1_PB_Q17, S2_ONT_Q17, S3_ONT_Q17, ONT_R9_longAIRR |
| " | spbc-error | 0.0 | All spatial datasets |
| " | spbc-failed | TRUE | All spatial datasets |
| " | spbc-cut | TRUE | All spatial datasets |
| " | spbc-length | 16 | S1_ONT_Q10, S1_ONT_Q17, S1_ONT_Q17_duplex, S1_PB_Q17, S2_ONT_Q17, S3_ONT_Q17, ONT_R9_longAIRR |
| " | group-field | SPBCUMI | S1_ONT_Q10, S1_ONT_Q17, S1_ONT_Q17_duplex, S1_PB_Q17, S2_ONT_Q17, S3_ONT_Q17, ONT_R9_longAIRR |
| " | group-field | UMISPBC | S4_ONT_Q17_HD |
| " | minl | 800 | All spatial datasets |
| " | maxl | 3800 | All spatial datasets |
| " | adaptive-filter | TRUE | All spatial datasets |
| " | filter-min-size | 5 | All spatial datasets |
| " | bin-size | 100 | All spatial datasets |
| " | peak-margin | 0.05 | All spatial datasets |
| " | cons-gap | 0.5 | All spatial datasets |
| " | cons-error | 0.2 | All spatial datasets |
| " | n-subsample | 750 | All spatial datasets |
| " | n-chunk | 75 | All spatial datasets |
| " | seed | 1 | All spatial datasets |
| longairr seqtag | window-length | 3000 | All spatial datasets |
| " | anchor-error | 0.1 | All spatial datasets |
| " | pf | ISOTYPE | All spatial datasets |
| " | split | Ig,TCR | S1_ONT_Q10, S1_ONT_Q17, S1_ONT_Q17_duplex, S1_PB_Q17, S2_ONT_Q17, S3_ONT_Q17, S4_ONT_Q17_HD |
| " | split | TCR | ONT_R9_longAIRR |
| " | split-regex | Ig TCR | S1_ONT_Q10, S1_ONT_Q17, S1_ONT_Q17_duplex, S1_PB_Q17, S2_ONT_Q17, S3_ONT_Q17, S4_ONT_Q17_HD |
| " | split-regex | TCR | ONT_R9_longAIRR |
| longairr airr | ph | TRUE | All spatial datasets |
| " | loci | ig | S1_ONT_Q17_duplex, S1_PB_Q17, S2_ONT_Q17, S3_ONT_Q17, S4_ONT_Q17_HD |
| " | loci | tr | All spatial datasets |
